# From individuals to populations: How intraspecific competition shapes thermal reaction norms

**DOI:** 10.1101/513739

**Authors:** François Mallard, Vincent Le Bourlot, Christie Le Coeur, Monique Avnaim, Romain Péronnet, David Claessen, Thomas Tully

**Affiliations:** Institut d’écologie et des sciences de l’environnement de Paris, CNRS - UMR 7618, iEES Paris, Sorbonne Université, 4 Place de Jussieu, 75005 Paris, France; Institut de Biologie de l′ENS (IBENS), Ecole Normale Supérieure, 46 rue d’Ulm, 75005 Paris, France; Department of Biology, Vesilinnantie 5, Natura 2nd floor, FI-20014 University of Turku, Finland

**Author notes:** deceased July 2018 (David Claessen).

**Keywords:** Body length, Collembola, Demography, Density-dependence, Growth Trajectory, Plasticity, Population Dynamic, Temperature Size Rule

## Abstract

1. Most ectotherms follow the temperature-size rule (TSR): in cold environments individuals grow slowly but reach a large asymptotic length. Intraspecific competition can induce plastic changes of growth rate and asymptotic length and competition may itself be modulated by temperature.
2. Our aim is to disentangle the joint effects of temperature and intraspecific competition on growth rate and asymptotic length.
3. We used two distinct clonal lineages of the Collembola *Folsomia candida*, to describe thermal reaction norms of growth rate, asymptotic length and reproduction over 6 temperatures between 6°C and 29°C. In parallel, we measured the long-term size-structure and dynamics of populations reared under the same temperatures to measure growth rates and asymptotic lengths in populations and to quantify the joint effects of competition and temperature on these traits.
4. We show that intraspecific competition modulates the temperature-size rule. In dense populations there is a direct negative effect of temperature on asymptotic length, but there is no temperature dependence of the growth rate, the dominant factor regulating growth being competition. We fail to demonstrate that the strength of competition varies with temperature except at the lowest temperature where competition is minimal. The two lineages responded differently to the joint effects of temperature and competition and these genetic differences have marked effects on population dynamics along our temperature gradient.
5. Our results reinforce the idea that the TSR response of ectotherms can be modulated by biotic and abiotic stressors when studied in non-optimal laboratory experiments. Untangling complex interactions between environment and demography will help understanding how size will respond to environmental change and how climate change may influence population dynamics.

## Introduction

(Angilletta, 2009; Atkinson, 1996; Edeline, Lacroix, Delire, Poulet, & Legendre, 2013; Gillooly, Brown, West, Savage, & Charnov, 2001). Thermal reaction norms of these traits have typically unimodal asymmetric shapes: the trait’s performance first increases more or less linearly with increasing temperature, reaches a maximum at some optimal temperature and then decreases rapidly above this optimum (Kingsolver, 2009). Thermal reaction norms of asymptotic body length differ: most ectotherms follow the temperature-size rule (TSR), which states that adult body length decreases with increasing temperature despite an increase in average growth rate (Zuo, Moses, West, Hou, & Brown, 2011; DeLong, 2012; Gardner, Peters, Kearney, Joseph, & Heinsohn, 2011; Walters & Hassall, 2006; Angilletta, 2009; Daufresne, Lengfellner, & Sommer, 2009; Atkinson, 1994). The TSR is defined within thermally favourable conditions (Atkinson, 1994), which is often viewed as the thermal range between a minimal and an optimal temperature (Walczyńska, Kiełbasa, & Sobczyk, 2016). Thus, most studies report measurements of reaction norms in optimal environments (including unlimited access to food resource) in order to avoid confounding indirect effects of temperature through density-dependence effects. Measurements are usually made on isolated individuals (Driessen, Ellers, & Van Straalen, 2007; Ellers & Driessen, 2011; Hoefnagel, de Vries, Jongejans, & Verberk, 2018), small cohorts (Karan, Morin, Moreteau, & David, 1998; Liefting, Hoffmann, & Ellers, 2009; Karan et al., 1998; Liefting et al., 2009; Ghosh, Testa, & Shingleton, 2013; Hoefnagel & Verberk, 2015) or growing populations with reduced density-dependence effects (Walczyńska et al., 2016) reared in the laboratory.

Natural insect populations are facing rapid temperature changes and there is need to understand and predict how populations will respond to these changes (Bewick, 2016). Yet, little is known on the robustness of the TSR predictions in a population context: as inter-individual interactions such competition might be thermally dependent, the picture becomes more complex. First, direct effects of temperature - such as the TSR - will affect demography: in ectotherms, warming accelerates life cycles by increasing growth rates, by speeding up maturation, but also by shortening lifespan (Kelly, Zieba, Buttemer, & Hulbert, 2013; Kingsolver & Huey, 2008). In parallel, warming can lower fecundity by reducing adult body length. Second, indirect effects of temperature mediated by demography will alter individual responses. For example, the intensity of competitive interactions may vary with temperature (Gherardi, Coignet, Souty-Grosset, Spigoli, & Aquiloni, 2013; Nilsson-Örtman, Stoks, & Johansson, 2014). Understanding the link between competition and temperature can thus become crucial in order to predict both individuals’ and populations’ responses to thermal variations. For example, studying the joint effects of temperature and intraspecific competition is required to understand the relative dominance of coexisting ants species (Cerdá, Retana, & Manzaneda, 1998; Diamond et al., 2017; Bestelmeyer, 2000). Population dynamics of a bordered plant bug are also only explained when taking into account the joint effect of temperature and food seasonal fluctuations together with density dependence effects (Johnson et al., 2015).

Intraspecific competition is one of the factors known to modulate populations’ responses to temperature variation (Bassar, Letcher, Nislow, & Whiteley, 2016). Intra-specific competition is often characterised as a density-dependent process (such as food limitation) that can lead to large variation in population sizes (Klomp, 1964). A recent study proposed two alternative hypotheses on the temperature-dependence of intraspecific competition in ectotherms (Amarasekare & Coutinho, 2014). First, the strength of intraspecific competition can increase monotonically with temperature. This is expected when resource requirements increase with temperature due to higher activity levels (Brown, Gillooly, Allen, Savage, & West, 2004; Ohlberger, Edeline, Vøllestad, Stenseth, & Claessen, 2011; Savage, Gillooly, Brown, West, & Charnov, 2004). In this scenario, direct and indirect effects of temperature act synergistically: above a physiologically optimal temperature, direct negative effects of warming will be amplified by a monotonically increasing strength of competition with increasing temperature. Alternatively, the strength of intraspecific competition can reach a maximum at intermediate temperature, near the optimal temperature for reproduction (such as in the example of the bordered plant bug in (Johnson et al., 2015)). The increase of re-source uptake required to maximize reproduction at intermediate temperatures strengthens the competition (Amarasekare & Coutinho, 2014). In this second scenario, direct and indirect effects will act antagonistically: above the thermal optimum, warming will have a direct negative effect but an indirect positive effect due to the loosening of the strength of competition with increasing temperature. This second scenario is expected to generate more complex demographic responses (Amarasekare & Coutinho, 2014). Note that for temperatures below the optimal temperature (where the TSR is defined), the two hypotheses have the same predictions as direct and indirect effects always acts antagonistically.

Our aim is to investigate how the effects of temperature on intraspecific competition affect the individual’s life history responses to temperature. We used the parthenogenetic Collembola *Folsomia candida* (Willem, 1902;) as an experimental model. We measured in parallel the responses of both isolated individuals and populations maintained at different temperatures. We compared two genetically distinct clonal lineages (labelled HA and TO) to study how within-species variations at individual level integrate with demography (Figure 1). These two lineages have two contrasted life history strategies along a slow (HA) - fast (TO) continuum (Mallard, Farina, & Tully, 2015; Tully & Ferrière, 2008) and may differ in their competitive abilities. Note that we only surveyed monoclonal populations. Thus competition only occurred between individuals sharing the same genotype. We did not study how the two lineages interact with each other.

**Figure 1:**
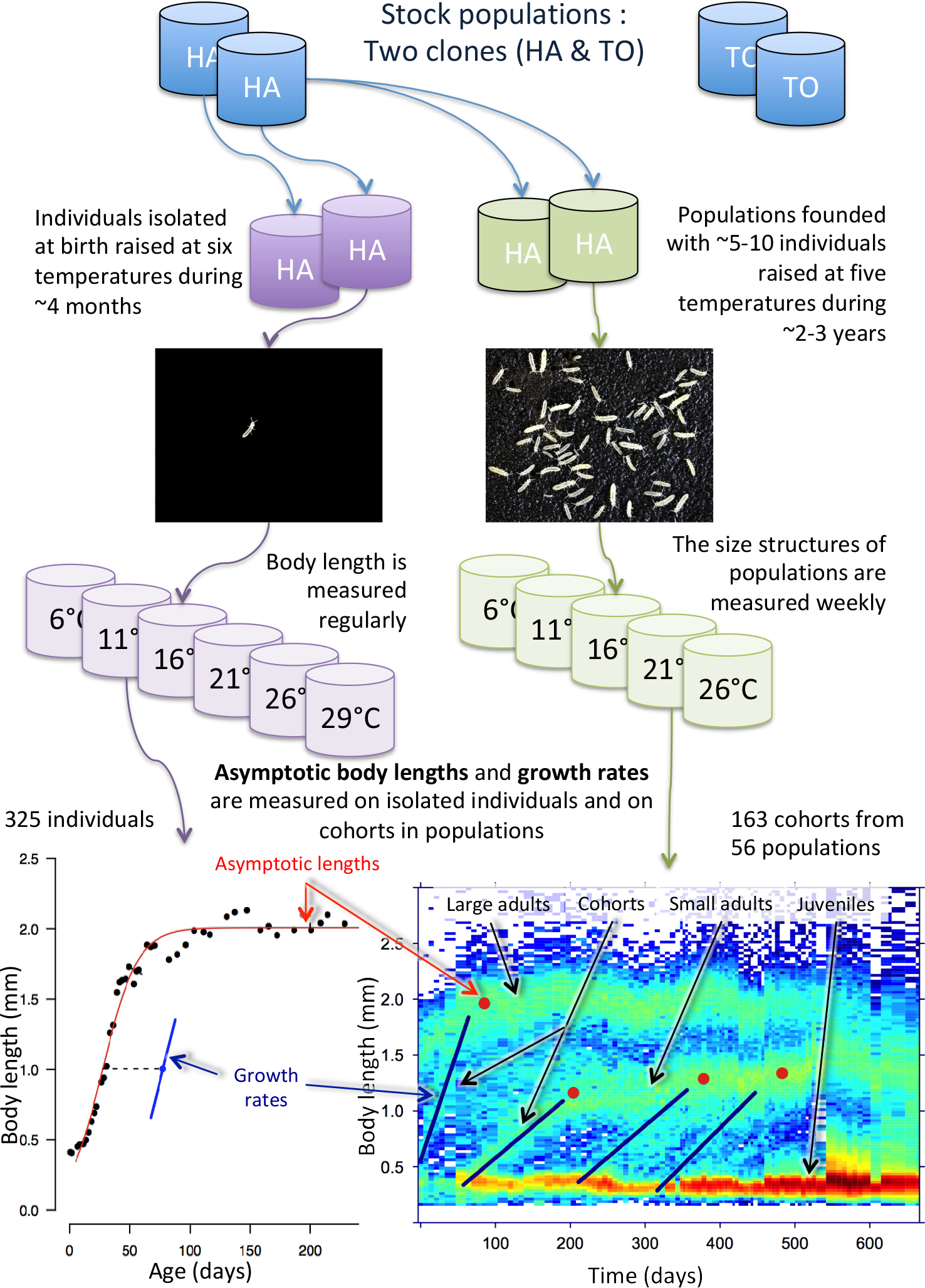
Experimental design. Two genetically different clonal lineages have been studied (HA & TO). Isolated individuals and populations have been maintained and studied at respectively six and five temperatures. Growth trajectories and long-term structure population dynamics (also called *structure time diagrams*) have been used to measure asymptotic body lengths and growth rates. Structure time diagrams display temporal dynamics of the populations’ size-structure: for each time (x-axis) and size class (y-axis) coordinate, a colour rectangle is plotted whose hue refers to the number of individuals on a log scale (warmer colours are higher abundance). Dark blue lines are examples of some measurements of cohort growth rates. Red dots are estimates of the mean asymptotic length attained by these cohorts. Note that in some populations such as the one displayed here (HA maintained at 21°C), the size structure can be trimodal with two modes of adults coexisting with the mode of juveniles. In such cases, adult cohorts whose mean asymptotic body lengths remain below 1.5 mm are referred to as “small adults”. See main text for details.

In this species, the strength of competition is expected to increase (i) with adult density because large individuals are known to monopolise most of the food resource through interference competition (Le Bourlot, Tully, & Claessen, 2014; Le Bourlot, 2014) and (ii) with the rate of reproduction which reflects the resource requirements (Tully & Ferrière, 2008). Based on (i) we can anticipate that the temperature induced variations in competition intensity will be linked to the effect of temperature on adult density in populations. Based on (ii) we can expect that the strength of competition will be maximal at temperatures that are optimal for reproduction, which may differ between the two lineages.

## Materials and methods

### Experimental setup

We used the Collembola *Folsomia candida* - a small (~2 mm long) blind ametabolous hexapod - as a model organism to run experiments on both isolated individuals and populations. Details on rearing conditions and lineages used in experiments can be found in the Supporting Information section. Briefly, we followed individuals from two clonal lineages (labelled TO and HA) either isolated or in populations at six different temperatures (6°, 11°, 16°, 21°, 26° and 29°C). Individuals were followed from hatching until death and size-structure of populations was measured weekly for more than a year. Populations were kept long enough (several months or years) to allow them to regulate themselves with density dependence. Competition between individuals is therefore intense in these monoclonal populations. Populations are self-regulated by the resource - delivered weekly - that is kept constant over time and between temperatures. This resource became progressively limiting in every populations (i.e. no remaining food after a week) except at 6°C.

### Measurements made on isolated individuals

#### Reproduction

Containers were regularly inspected for eggs in order to count the number of clutches and eggs laid by each individual (n=185) during the experiment (400 days). The total number of eggs laid by each individual was used as an overall measurement to determine optimal temperatures for reproduction (Figure 2A).

**Figure 2:**
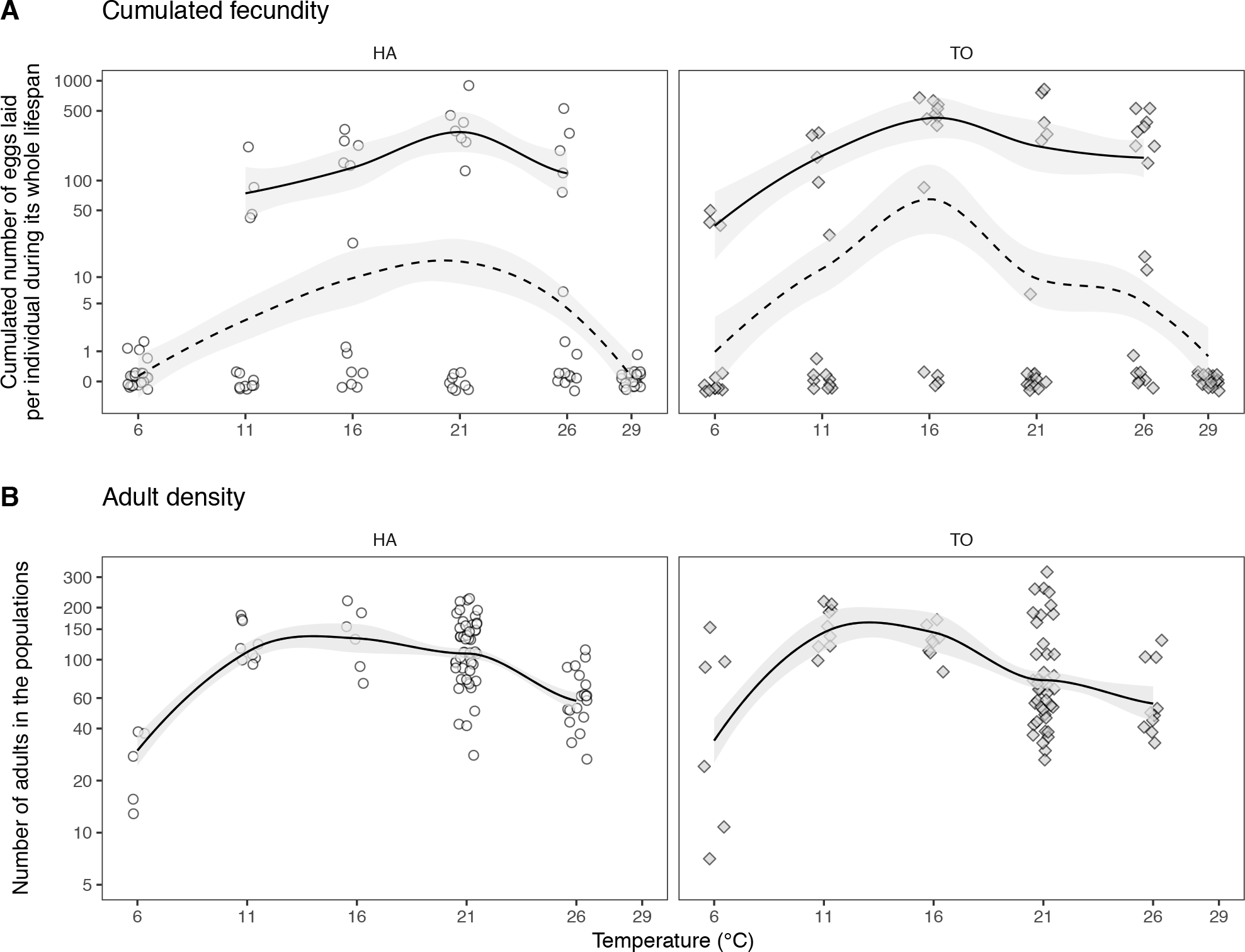
The effects of temperature on reproduction of isolated individuals (A) and on density in populations (B) for lineages HA (open circle) and TO (grey diamonds). The panel A represents the total number of eggs laid by an individual during its whole life. The strait line is a smooth function fitted on the reproductive adults only excluding individuals that died before reproducing (fecundity higher than zero), while the dotted line is adjusted on the whole dataset which includes individuals that did not reproduce. For both measurements, the maximal reproduction peaks around 21°C for HA and around 16°C for TO. The panel B displays adult densities measured in populations at the time of measurements of growth and asymptotic length (see methods for details). On average maximum adult densities are reached between 11°C and 21 °C.

#### Growth rate and asymptotic body length of isolated individuals

During the first ten weeks of the experiment, individuals were measured three times per week. Then, after ten weeks, when their growth has slowed down, they were measured once a week. This was done until all individuals died (400 days, 95 % of the individuals died before 230 days). For each body length measurement, two to three pictures were taken using a digital camera (Nikon D300) fixed on a dissecting scope. Body length was measured on each picture using the ImageJ software (Abramoff, Magalhaes, & Ram, 2004) (http://rsbweb.nih.gov/ĳ/). More precisely, we measured the length from the front of the head to the rear of the abdomen on pictures taken from above such as the one visible on Figure 1. This measurement was done using the “segmented line selection tool”, after calibrating units with the “set scale” command. Smoothed splines growth curves were fitted to each individual growth trajectory (Figure 1) using the function *gcFitSpline* from the package *grofit* with a Gompertz type function. We adjusted the smoothness of the spline fit so as to generate estimates of growth parameters that are comparable to the ones measured on populations (see Figure S1 for details). Growth rates and asymptotic lengths were estimated from these fits (parameters *mu* and *A*).

### Measurements made on cohorts in populations

#### Dynamics of the population size structure

The population size structure was measured weekly during more than a year using a dedicated program. Each measurement gave us the number of individuals and, for each individual counted, its body length (mm). Given the number of Collembola in a box, and that individual marking is unattainable, it was not possible to measure the asymptotic body length and growth rate of an individual in a population. But we managed to extract measurements of these life history traits in populations using the graphical representation for structured time series (Figure 1 and Figure S2) provided by the R package *STdiag* (Le Bourlot, Mallard, Claessen, & Tully, 2015). The examination of the structure time diagrams reveals many breakaway waves of density through time in populations (Figure 1). These waves of density - which we call “cohorts” - connect small to large individuals over time and result from the synchronised growth of a group of small Collembola within the group of larger individuals (See the “Cohort” in the bottom right panel of Figure 1 and for example the TO_06_r2 and TO_06_r4 populations in Figure S2). We visually inspected the structure time diagrams of our 56 populations to identify 163 cohorts that are sufficiently contrasted graphically to be studied quantitatively (Figure 1, Figure S2, Figure S3). Note that we did not take into account juveniles that, for different reasons, never managed to grow, or those that did not grow in a cohort and whose growth trajectories were thus invisible on graphs.

#### Mean growth rate and asymptotic body length in populations

We measured the mean growth rate and asymptotic body length of the 163 distinct cohorts that we identified. The black segments on the structure-time diagram of Figure 1 (bottom right panel) shows how we measured the mean growth rate in the populations: it is the slope of a line adjusted through visible growing cohorts (waves). The asymptotic body length was estimated visually as the mode length of individuals once the growing cohort had fully merged with pre-existing adults, or when the mode length of the growing cohort stabilizes (black dots on the structure-time diagram of Figure 1). Note that in some HA populations at 21°C, two modes of large individuals could be identified: some of the measured cohorts produced large fully-grown individuals (>1.5 mm) while others produced individuals that stabilised their growth at smaller sizes (<1.5 mm, Figure S2 and Figure 3B). More details on these measurements are provided in Supporting Information section and in Figure S1.

**Figure 3:**
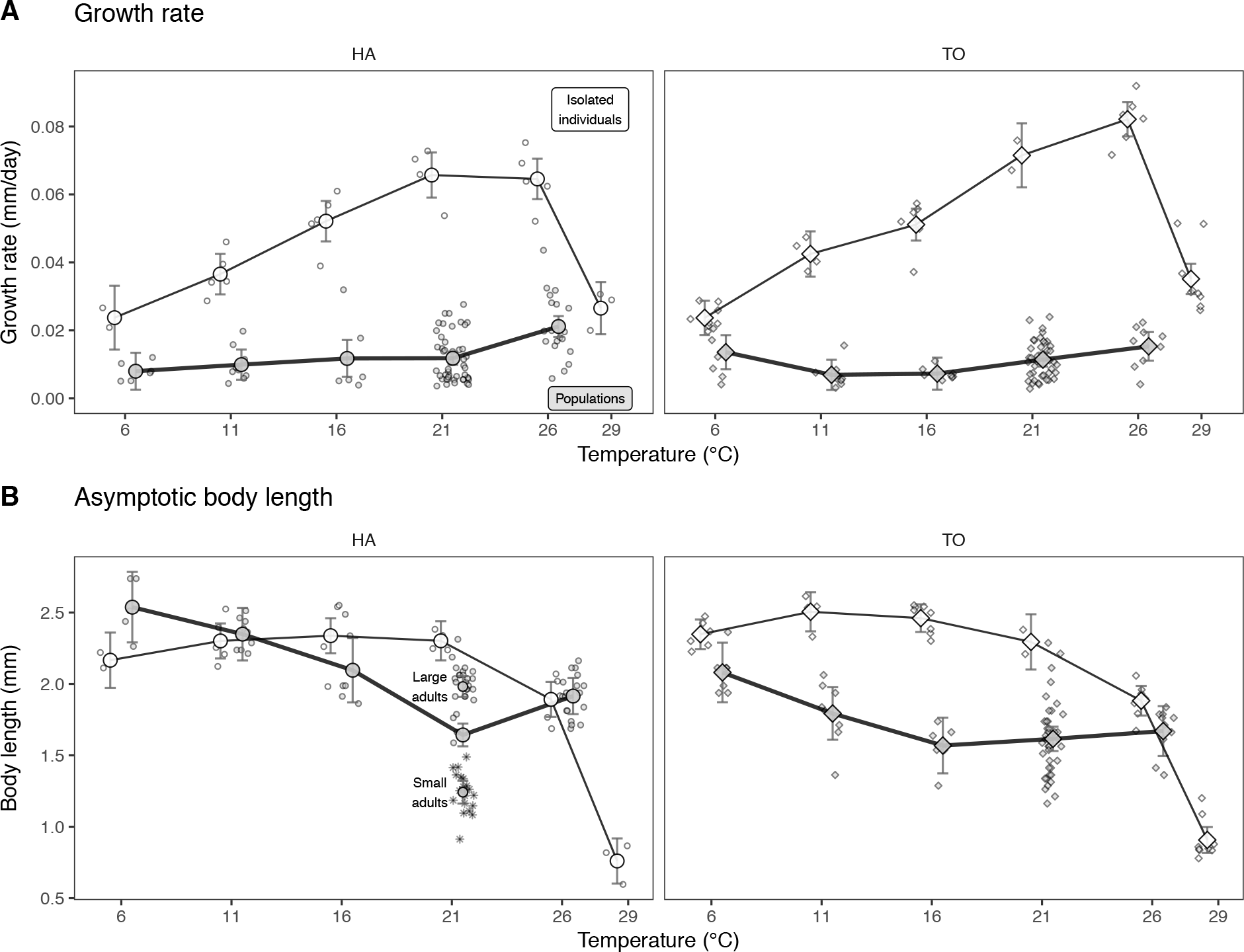
The thermal reaction norms of growth rate (A) and asymptotic body length (B) for lineages HA (left, circles) and TO (right, diamonds). Means (and 95% confidence intervals) are plotted for each combination of lineage identity, temperature and type of measurement (isolated individuals and cohorts in populations) together with the raw measurements. Open symbols and fine line are used for measurements made on isolated individuals and symbols filled with grey and thick solid lines are used for measurements made in populations. Asterisks are used to represent small adults observed in some HA populations at 21°C. Note that at 29°C we only plot measurements made on isolated individuals because we failed at maintaining populations at 29°C since individuals do not reproduce at this temperature (Figure 2A).

#### Measuring densities of large and small individuals

Using the R package *STdiag*, we also extracted mean densities of adults and juveniles. More technical details can be found elsewhere (Le Bourlot et al., 2015) and in Supporting Information. Individuals are considered as juveniles if they are shorter than 0.3 mm and as large if they are longer than 0.9 mm based on the multimodal body length distributions in the populations (Figure S6). We use the term “adult” to designate large individuals in populations although it does not mean that they are all reproductive adults since we cannot measure the size at maturity in populations.

### Measuring joint effects of temperature and intraspecific competition in populations

In populations, changes in growth rates and asymptotic body lengths are expected to be determined by direct effects of temperature (warming increases growth rate and downsizes body length) and by direct effects of intraspecific competition (density dependence). But temperature can also intervene through indirect effects by modifying the population density and the strength of intraspecific competition.

To study whether indirect effects of temperature (via competition) alter patterns seen at the individual level, we first compared isolated individuals with cohorts, without considering the effect of density within populations (Figure 3, Table 1, Table 2). We then took advantage of long-term variations of adult densities within populations (Figure 2B & Figure S2) to quantify the strength of competition for each combination of temperature and lineage (Figure 4, Table S1) and to study if, and how, the strength of competition changes with temperature (Figure 5A, B). The strength of intraspecific competition was estimated as the effect of the logarithm of adult density on the growth rate and asymptotic length (Table S1, Figure 2B).

**Figure 4:**
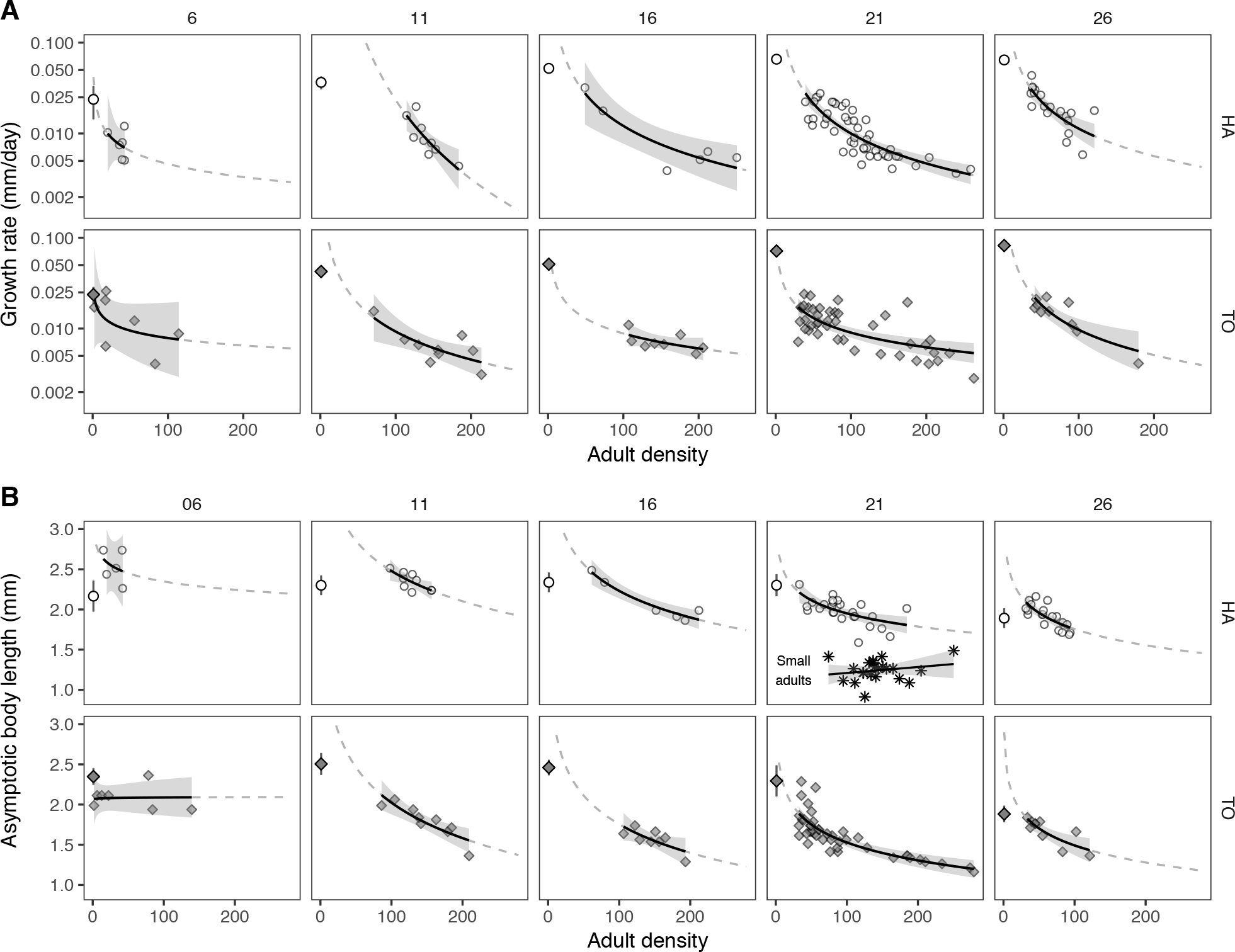
The joint effects of temperature (four columns of panels) and adult density (mean number of adults per container) on cohort growth rates (A) and asymptotic body length in populations (B) for the two lineages HA (open circles) and TO (filled diamonds). On the left side of each panel the mean growth rate and asymptotic body lengths (95%CI) measured on isolated individuals (Figure 4 A and B) are plotted for comparisons. Growth rates are plotted on a log10 scale. The function adjusted to the data is a linear fit of the form y=a+b*log(x), x being the mean density of large adults at the time of the measurement. It is prolonged out of the range of the data with a dotted curve. The size of the “small” adults observed in the HA populations at 21°C are plotted with asterisks.

**Figure 5:**
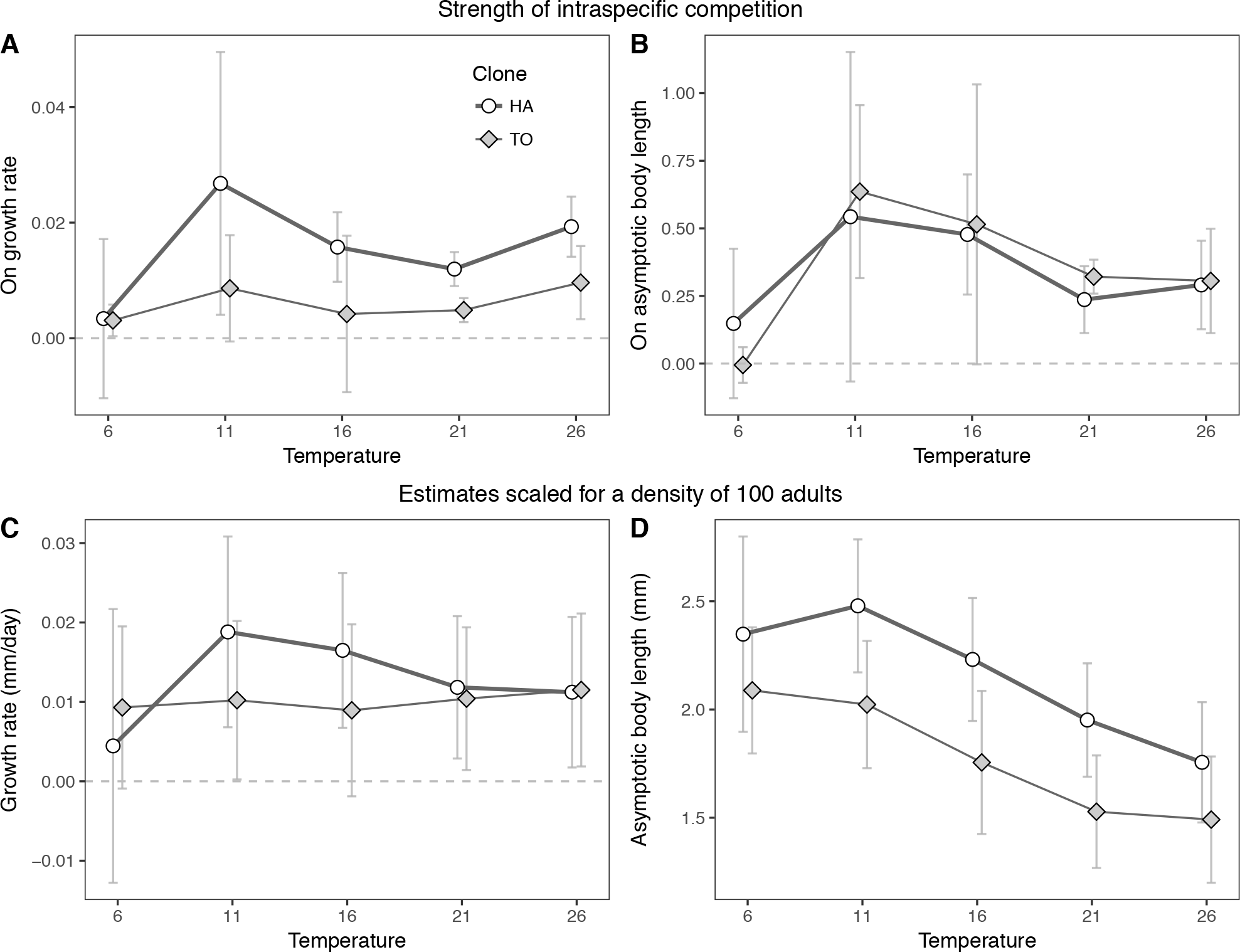
The strength of intraspecific competition within populations has been estimated as the slope (reversed to positive values) of the effect of adult density (log scale) on the growth rate (A) or asymptotic body length (B) estimated independently here for each lineage and temperature (mean ± 95% confidence intervals). To visually reveal the effect of temperature on the growth rate (C) and asymptotic body length (D) in populations while controlling for density we have plotted the predicted estimates of growth rate and body length for an adult density scaled at 100 individuals per box using an unconstrained linear model with an interaction between lineage and temperature.

**Table 1:** The two models of this table have been done to study whether indirect effects of temperature (via competition) alter patterns seen at the individual level. Model A (growth rate) is associated with the Figure 3A and model B (asymptotic length) with Figure 3B. In both models, the variable *lineage* has two levels (HA, TO), *temperature* has 5 levels (29°C is not included since we have no population at this temperature) and the variable *competition* is used to tell the difference between measurements made on isolated individuals with the ones made on populations. Non-significant interactions that have been dropped from initial full models are shown in italics. Effects and their interactions are tested with likelihood ratio tests using a type 3 anova (*Anova* function from the *car* package). Parameters of full models A and B are provided in Table S3.

| Factors | Likelihood ratio Chi-square | Df | P (>Chisq) |
| --- | --- | --- | --- |
| **(A) Growth rate** |  |  |  |
| Lineage (HA/TO) | 2.3 | 1 | 0.13 |
| Temperature (factor) | 35.9 | 4 | <0.001 |
| Competition (isolated/population) | 8.5 | 1 | 0.003 |
| Temperature*Competition | 50.6 | 4 | <0.001 |
| Lineage*Competition | 0.76 | 1 | 0.38 |
| Lineage*Temperature | 8.1 | 4 | 0.09 |
| Temperature*Lineage*Competition | 16.1 | 4 | 0.0028 |
| **Growth rate for HA** |  |  |  |
| Temperature (factor) | 27.2 | 4 | <0.001 |
| Competition (isolated > population) | 6.5 | 1 | 0.011 |
| Temperature*Competition | 38.3 | 4 | <0.001 |
| **Growth rate for TO** |  |  |  |
| Temperature (factor) | 17.2 | A | 0.002 |
| Competition (isolated > population) | 12.4 | 1 | <0.001 |
| Temperature*Competition | 237.1 | 4 | <0.001 |
| **(B) Asymptotic body length** |  |  |  |
| Lineage (HA/TO) | 9.4 | 1 | <0.001 |
| Temperature (factor) | 99.4 | 4 | <0.001 |
| Competition (isolated/population) | 3.1 | 1 | 0.09 |
| Temperature*Competition | 35.0 | 4 | <0.001 |
| Lineage*Competition | 22.2 | 1 | <0.001 |
| Lineage*Temperature | 19.6 | 4 | <0.001 |
| *Temperature\*Lineage\*Competition* | 8.6 | 4 | 0.07 |
| **Asymptotic body length for HA** |  |  |  |
| Temperature (factor) | 78.3 | 4 | <0.001 |
| Competition (isolated~population) | 2.1 | 1 | 0.14 |
| Temperature*Competition | 17.8 | 4 | 0.001 |
| **Asymptotic body length for TO** |  |  |  |
| Temperature (factor) | 37.7 | 4 | <0.001 |
| Competition (isolated > population) | 6.2 | 1 | 0.012 |
| Temperature*Competition | 32.8 | 4 | <0.001 |

**Table 2:** This table represents the best selected linear models that we used to study effects of lineage (HA and TO) and temperature (as a factor) on growth rate (A, B, Figure 3A) and asymptotic size (C, D, Figure 3B) measured either on isolated individuals (A, C, Figure 3 open symbols and thin line) or in populations (B, D, Figure 3, filled symbols and thick lines). Small and large HA adults at 21°C are taken into account in the model. Interactions that have been dropped from initial full models are italicized. Effects and their interactions are tested with likelihood ratio tests using a type 3 anova (*Anova* function from the *car* package). We also report the effects of temperature tested independently for the two clones for models A and D since the interaction between lineage and temperature was significant.

| Factors | Likelihood ratio Chi-square | Df | P (>Chisq) |
| --- | --- | --- | --- |
| **(A) Growth rate of isolated individuals** |  |  |  |
| Lineage (HA/TO) | 0.00 | 1 | 0.99 |
| Temperature (factor) | 116.6 | 5 | <0.001 |
| Lineage*Temperature (6-16°C:HA=TO; 21-29°C:HA<TO) | 11.9 | 5 | 0.035 |
| Temperature (HA only) | 102 | 5 | <0.001 |
| Temperature (TO only) | 322 | 5 | <0.001 |
| **(B) Growth rate in populations** |  |  |  |
| Lineage (HA=TO) | 2.3 | 1 | 0.13 |
| Temperature (factor) | 35.8 | 4 | <0.001 |
| *Lineage\*Temperature* | 8.08 | 4 | 0.089 |
| **(C) Asymptotic body length of isolated individuals** |  |  |  |
| Lineage (HA<TO) | 8.0 | 1 | 0.0047 |
| Temperature (factor) | 1105 | 5 | <0.001 |
| *Lineage\*Temperature* | 4.9 | 5 | 0.42 |
| **(D) Asymptotic body length of cohorts in populations** |  |  |  |
| Lineage (HA>TO) | 7.8 | 1 | 0.005 |
| Temperature (factor) | 91 | 4 | <0.001 |
| Lineage*Temperature | 23.2 | 4 | <0.001 |
| Temperature (HA only) | 69.0 | 4 | <0.001 |
| Temperature (TO only) | 30.6 | 4 | <0.001 |

To quantify the direct effects of temperature in populations, we controlled for density by scaling all populations as if adult densities in containers was constant and equal to 100 adults (Figure 5C, D). This corresponds to a density observed in almost every combination of lineages and temperatures (Figure 2B) and is close to the overall mean adult density observed in the populations (overall mean: 103 adults per container).

Lastly, to better visualize the joint effects of temperature and density on the two traits, we grouped the measurements made on isolated individuals (“adult density” of zero) and those made in populations at the different adult densities to draw a 3d contour plot using the function *stat_contour* from the *ggplot2* library (Figure S7).

### Statistical analysis

All analyses were done with R 3.3.1 software http://cran.r-project.org (Ihaka & Gentleman, 1996) and we used the package *ggplot2* to produce the graphics (Wickham, 2009).

#### Statistical models

We used Gaussian linear models (*glm* function) to study the growth rate and asymptotic length. We started with full models with all dependent variables and their interactions and simplified these models by removing non-significant interactions. The factors lineage (the two clones) and temperature were included in the initial models. The effects and their interactions are tested with type 3 anova (*Anova* function from the package *car*). In statistical tables (Table 1, Table 2, Table S1) we report in italics the tests corresponding to complex interactions that have been dropped from the full models and results of the type 3 anova on simplified models. We performed several complementary analyses on subsets of the whole datasets to better understand complex interactions and to perform specific comparisons. For instance, when we found one or several significant interactions between lineages and other variables, we performed some independent sub-analyses on each genetic line to better understand each of their specific responses.

#### Temperature

Temperature often had nonlinear effects. Thus in the statistical models we generally considered temperature as a categorical variable with 6 levels for isolated individuals and 5 levels for the populations (29°C missing) to avoid imposing a priori a parametric response. In some models, or for some specific comparisons, we sometimes considered the temperature as a continuous variable when the response to temperature was visually linear (between 6° and 26°C for the growth rate of isolated individuals for instance, Figure 3A, B).

#### Graphics

We do not report values of estimates from the models in the tables but we refer to the graphics that have been designed to help understand and support the statistical analysis. Means and their 95% confidence intervals on figures, and log-linear relationships and their 95% confidence intervals in Figure 4 were estimated using saturated models including all the complex interactions. We used these saturated models to produce figures that reveal the data with minimal constrains (Tufte, 2001). Estimates and 95%confidence intervals of parameters of the saturated models are provided in Table S3.

## Results

### Temperature, reproduction and adult density

#### Reproduction

Cumulative reproduction of isolated individuals is reduced at the lowest temperature extreme (6°C) and is absent at the highest one (29°C) for both lineages. At 6°C, growth is so slow that it takes more than 100 days for most individuals to reach maturity. At 29°C, the reproductive cessation could result from the asymptotic body lengths of individuals (≤1mm, see below) being smaller than the average size at maturity at 29°C (~1.2-1.4mm) (van Dooren, Tully, & Ferrière, 2005). Between these extremes, the mean cumulative reproduction has an unimodal bell-shaped thermal response curve with a maximum reproduction reached at a lower temperature for TO (~16°C) than for HA (~21°C, Figure 2A). TO has on average a higher fecundity than HA (Figure 2A, Figure S4A). For both lineages, the mean clutch size measured on the first 5 clutches decreases with increasing temperature but TO lay larger clutches than HA (Figure S4A). On average, at low temperature, springtails lay larger clutches but less often than at warmer temperatures.

#### Adult and juvenile density

We counted on average 100 large individuals per population but this number ranged from less than 20 to more than 200 because of long-term fluctuations in populations′ densities and structures (Figure 2B, Figure S2). The average adult density is low at the two temperature extremes, and reaches a maximum between 11° and 21°C (Figure 2B). Adult densities do not significantly differ between lineages. The density of juveniles is also variable within and across treatments. Below 21°C, TO populations bear on average more juveniles than HA (Figure S4B).

### The effect of temperature on growth rate

#### Individuals versus populations

A full model combining measurements made on isolated individuals and on cohorts shows that the growth rate is influenced by a complex interaction between lineage identity, temperature and competition treatment (isolated individuals versus population) and by an interaction between temperature and competition (Table 1A, Figure 3A): isolated individuals grew on average 4.2 time faster than cohorts in populations on the whole temperature range but the difference between the two competition treatments was stronger for intermediate temperatures (5.6 times faster at 16°C and 5.9 times faster at 26°C, Figure 3A). The average effect of temperature on growth rates of isolated individuals (+2.74 μm/day/°C, average estimated over the two lineages) is 8.7 times higher that the mean effect of temperature on growth rates in populations (+0.31 μm/day/°C, Figure 3A). We further split the data to examine the effects of temperature and lineage identity in each competition treatment separately (isolated individuals/ populations).

#### Individuals

Growth rates of isolated individuals depend on an interaction between lineage identity and temperature but, as expected, the effect of temperature was massive for both lineages (Table 2A). Growth rates increase almost linearly with temperature (Figure 3A) between 6° and 26°C (2.56 ± SE. 0002 μm.day^−1^.°C^-1^, 95% CI: 2.22-2.29). They then drop between 26° and 29°C. Both lineages have similar growth rates on the whole temperature range except at 26°C where TO grows on average 30% faster than HA (t=4.06, p = 0.0023).

#### Populations

Growth rates measured in populations did not differ between clonal lineages but varied slightly with temperature (Table 2B): on average, cohorts are growing 75% faster at 26°C than at the colder temperatures combined (contrast between 26°C and the four other temperatures F_1,162_ = 35.7, p < 0.001) while the growth is uniformly low over the four lower temperatures (F_3,131_ = 1.7, p = 0.17).

### The effect of temperature on asymptotic body length

#### Individuals versus populations

Using the whole dataset, we found that the asymptotic body length is affected by significant two-way interactions between competition (isolated individuals versus populations), lineage identity and temperature (Table 1B).

For HA, the asymptotic body length in population is very close to the length reached by isolated individuals except at 21°C where on average the small and large adults in populations are smaller than the isolated ones (Figure 3B, large adults χ^2^_1_=16, p < 0.001). Thus, for HA, 21°C is the only tested temperature where the asymptotic length in populations is smaller than the asymptotic length of isolated individuals (−0.66mm ± SE = 0.20 in populations, χ^2^_1_=10,7, p = 0.001).

Contrary to what was observed for HA, the asymptotic body length of TO in populations is significantly smaller than the asymptotic length of isolated individuals on the whole temperature range (comparisons for the five temperatures, χ^2^_1_> 8.4, p < 0.04). The difference between the two competition treatments is maximum at 16°C (−0.9mm ± SE = 0.06 in populations, χ^2^_1_ = 233, p < 0.001, Figure 3B).

#### Individuals

Lineage identity and temperature have additive effects on the asymptotic body length (Table 2C). On average isolated TO manage to reach a slightly larger asymptotic length than HA over the whole temperature range (+0.105 mm± SE = 0.037, χ^2^_1_=8.0, p = 0.005, Figure 3B). For the two clonal lineages, the asymptotic length of isolated individuals decreases non-linearly with temperature: the average length remains stable between 6° and 21°C (χ^2^_1_ = 1.07, p = 0.30) and then decreases abruptly between 21° and 29°C (Figure 3B) which is outside the optimal thermal range given that growth no longer increases with increasing temperature above ~21-26°C (Figure 3A). Thus, given that the temperature size rule is expected to occur for non-stressful temperatures (Atkinson, 1994; Walczyńska et al., 2016; Hoefnagel et al., 2018), isolated individuals appear to slightly or even not really follow the TSR, especially HA since its mean asymptotic lengths remain roughly stable on its non-stressful thermal range.

#### Populations

Lineage and temperature have additive effects on the mean asymptotic length in populations (Table 2D). Yet, HA that is on average larger than TO over the whole range of temperatures (+0.4 mm ± SE = 0.03, Figure 3B). The mean asymptotic length declines regularly between 6° and 21°C for HA and between 6° and 16°C for TO (~−0.048 mm.°C^−1^). Above these temperatures it either increases slightly (HA) or remains roughly constant (TO).

### The joint effects of temperature and density in populations

#### Growth rates

The full model that combines the two lineages shows that growth rates depend on two-ways interactions between lineage, temperature and adult density (Table S1 A). We further split the dataset to analyse separately the response of the two lineages (Table 1 B, C). We found that for both lineages growth rates of cohorts were determined by an additive effect of the adult density and the factor temperature (Table S1 B, C), which means that intraspecific competition (adult density on a log scale) had on average the same negative effect on growth rates for each temperature (Figure 4 A & Figure 5 A). For both lineages, the additive effect of temperature vanishes for temperatures above 6°C (Table S1 B, C). To better understand the biological meaning of the additive effect of temperature in populations for the two lineages (and its interaction with lineage identity in the main model Table S1 A), we plotted the predicted growth rates for the different temperatures in a virtual population where the adult density is fixed to 100 adults (Figure 5 C). This shows that, at this density, the growth rate is not influenced by temperature for TO but tends to decrease with increasing temperatures between 11° and 26°C for HA, although this change is not significant given the overlaps of the confidence intervals. Thus we conclude that at this adult density (100 ind.) the temperature has very limited direct effect on growth rates, which are mainly determined by the strength of intraspecific competition.

#### Asymptotic body length

In the full model that combines the whole dataset, the asymptotic body length in populations is affected by two significant interactions, *temperature*density* and *lineage identity*temperature* (Table S1 D). The first interaction means that the strength of intraspecific competition changes with temperature, similarly for the two lineages. This is visible on the Figure 5 B, which reports estimated values of an unconstrained fully saturated model (the one with all possible interactions). Competition is minimal at 6°C (see also the flat relation in Figure 4 B at 6°C) and maximal at 11°C. It then decreases progressively as the temperature increases.

As for the growth rates, we used the full model to predict - and plot - the asymptotic body length at the different temperatures in a population comprising 100 large individuals (Figure 5 D). This figure shows that the asymptotic body length decreases as temperature increases when the effect of density is controlled (−0.024mm/°C ± SE = 0.002 on average between 6° and 26°C) and that TO is on average smaller than HA (−0.41mm ± SE = 0.03) over the temperature range. The significant interaction between temperature and adult density for TO (Table S1 F) is due to the better estimation of the strength of competition at 6°C compared to HA (Table S1 E) for which we had fewer cohorts in populations raised at 6°C to estimate the asymptotic length.

### Analysing the contour plot

Lastly, the Figure S7 combines all the measurements that we made on isolated individuals and in populations to better understand visually the joint and non-linear effects of competition (density) and temperature. For TO, the intraspecific competition has a negative effect on asymptotic length with an additive effect of temperature that remains on almost the whole range of conditions explored. For HA, the density has first a positive effect on asymptotic length: when the adult density increases, the collembola tend to reach a longer asymptotic body length. But when the density continues to increase (above 50 individuals per container), its effect becomes negative.

Note that when comparing isolated individuals to populations we found that asymptotic lengths of HA are marginally modified in populations (Figure 3B), which seems to be in contradiction with the observed effect of intraspecific competition on this trait (Figure 5B). This is because density has first a positive effect on body length in this lineage (adults in small populations are larger than isolated individuals). This non-linear effect of density (visible on Figure 3B and Figure S7B) underlines the advantage of quantifying the strength of intraspecific competition by comparing populations with different densities rather than relying simply on the comparison of populations with isolated individuals since this would have led to conclude that competition has no effect on this trait.

## Discussion

As observed on many other ectotherms (Angilletta, 2009; Atkinson, 1996; Kingsolver & Woods, 1997), warmer conditions allow for faster collembola growth (Figure 3A, Figure 6) probably due to faster enzyme kinetics and increased food consumption (Angilletta, 2009). Despite this growth rate increase, the TSR only faintly applies: the isolated individuals maintained a large body length over a broad temperature range. A marked decrease in asymptotic body length was only detected above the optimal temperature range, where growth rates slow down and reproduction declines. Interestingly, in populations, the adult lengths decrease with increasing temperatures on the thermal range below the stressful warm temperatures (Figure 3B). This decrease is observed in both lineages and is even stronger when controlling for adult densities (Figure 5D). The decline in body length (−0.034mm/°C on average which corresponds to ~-1.7% reduction in body length per °C increase, Figure 5D) is within the range of values documented for terrestrial arthropods (Horne, Hirst, & Atkinson, 2015). Here, we only report asymptotic body length and the TSR might still apply to the size at maturity of *Folsomia candida* as it does for other Collembola species (Johnsen, 2014). Indeed, Hoefnagel et al. (Hoefnagel et al., 2018) have shown that, in *Daphnia*, different temperature-dependent mechanisms are explaining the TSR for maturity and asymptotic body length. They show that growth stops at smaller sizes in warmer environments because resource limitation becomes stronger at higher temperature and for larger body sizes. Since food availability is the main factor limiting population growth in our experiments, it is also likely that the same reason triggers the TSR in our populations while we almost did not detect it for isolated individuals. Above 26°C, isolated individuals arrest growth much earlier: there was no difference between isolated individuals and adults in populations. This early growth arrest could result from secondary constraints such as thermal instability or from an abrupt decline of the assimilation efficiency over ~21-26°C (Kukal & Dawson, 1989; McConnachie & Alexander, 2004). It has also been suggested that increased mortality rates above the thermal optimum could act as a selection pressure to decrease size at maturity (Kozlowski, 2004). The observed temperature-induced length plasticity may also be adaptive if the large individuals are more sensitive to heat stress than smaller ones. More generally, a reduction in asymptotic body length is likely a general response to stressful environments in arthropods with continuous growth such as *Folsomia candida* or *Daphnia magna* (Hoefnagel & Verberk, 2015). Some population structure-time diagrams also suggest that individuals can resume growth when the environmental conditions get better (a decrease in population density see Figure S2 population “HA_21_r1” around day 200) and can also shrink if the environment deteriorates (data not shown). Further experiments are required to fully characterise the flexibilities of growth and asymptotic size and to determine their adaptive values.

**Figure 6:**
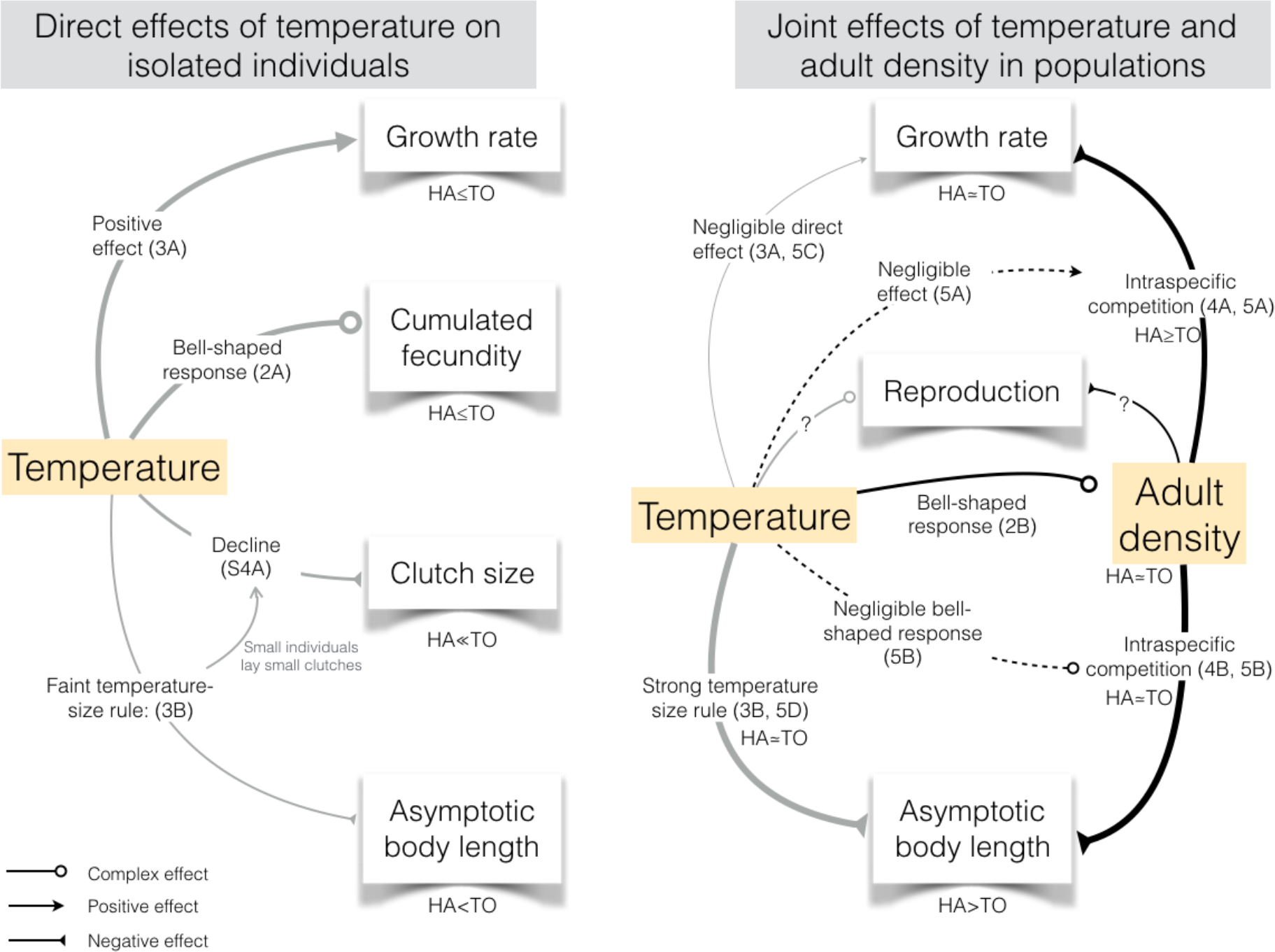
The direct and indirect effects of temperature on growth rates and asymptotic body lengths of isolated individuals (left) and of cohorts of individuals within populations (right). References in parenthesis point to figures. The line thickness underlines the relative strength of each effect. For example, adult density has a strong direct negative effect on both growth rate and asymptotic body length while temperature has a negligible direct effect on growth. Dotted lines represent the effects of temperature on the strength of intraspecific competition. Grey lines represent direct effects of temperature on life-history traits and black lines the effects linked with density in populations. Note that reproduction could not be measured in populations thus the probable effects of density and temperature on reproduction are mentioned with a question mark on the right side of the figure.

In the introduction, we discuss two scenarios for the temperature effect on intraspecific competition. Our results suggest that competition intensity is maximal at intermediate temperatures (Figure 2, Figure 5B) thus explaining differences between our lineages in their response to temperature in populations. As predicted by theory (Amarasekare & Coutinho, 2014), we observed that the temperatures at which TO and HA’s reproduction is maximal (16°C and 21°C, Figure 2A) are also the ones where the two lineages reach their minimal asymptotic body lengths in populations (Figure 3B). The existence of the trimodal adult size distributions at 21°C (Figure 2A) could also be interpreted as complex population dynamics under increased competition (Amarasekare & Coutinho, 2014), thus reinforcing our conclusion. After the establishment of a dense cohort of large adults, a second cohort can establish but at a smaller mean asymptotic length (see for instance population HA_21_11-16 in Figure S2). We have previously shown that large adults dominate smaller ones through intense interference competition (Le Bourlot et al., 2014) preventing them from growing further. A similar coexistence of different size classes of individuals has been shown in the lake perch (Persson et al., 2003). In this example, the population is alternatively dominated by stunted or giant individuals due to size-dependent cannibalism and intercohort competition. Here, our trimodal structures seem to persist as long as the dominant large adults remain numerous enough but may only establish in a new colonised environment with initially reduced competition for food. Finally, our measurements failed to detect any significant variation of the strength of competition on growth between 11° and 26°C (Figure 5A, B) as opposed to our findings on asymptotic length. Given the large confidence intervals (Figure 5), this discrepancy may be due to a lack of power to detect such a pattern. Indeed, the thermal responses shown in Figure 5B tend to support the idea that strength of competition is maximal at intermediate temperatures, despite large overlaps in confidence intervals. A similar result was described in the bordered plant bug, the effect of competition on fecundity was strongest at temperature optimal for reproduction (Johnson et al., 2015).

Overall, our study supports the idea that resource limitation is stronger for large individuals and at higher temperatures. At lower temperature, Hoefnagel et al. suggest that *Daphnia* might reach a “ceiling” asymptotic size that cannot be exceeded (Hoefnagel et al., 2018). While this might be the case for TO, we found that isolated HA individuals in optimal environments did not reach their highest possible size since larger asymptotic lengths are reached in populations. Assuming that the length reached by isolated individuals is optimal, exceeding this size will be associated with fitness costs. Conversely, these costs could be compensated by the advantageous of being among the largest individuals in populations. Under strong interference competition, giant individuals are advantaged to access the food (Le Bourlot et al., 2014). Further theory is required to interpret our observations as we only observed this increase in length at the lowest temperatures where densities and interference competition are low. Alternatively, the hoarding of nutritive resources by large adults in populations could serve as an ‘insurance’ to better face potential future degradation of the environment (Cuthill, Maddocks, Weall, & Jones, 2000) due for instance to resource depletion triggered by increased local density.

Finally, our results reveal genetic differences between our clonal lineages in their mean trait values and plastic responses to both temperature and intra-specific competition (Figure 6). When isolated, TO grow on average faster than HA in the hottest environments (>=21°C) and reach a larger body length (Figure 3). This difference is referred as a vertical or “faster-slower” shift: a change in the overall height of the thermal performance curves. This vertical shift is opposed to an horizontal or “hotter-cooler” shift: the variation in the location of the thermal maximum or in the position of the curve along the temperature axis (Kingsolver, 2009). While previous studies have described similar effects of temperature on Collembola growth rates (Birkemoe & Leinaas, 2000; Driessen et al., 2007; Ellers, Mariën, Driessen, & van Straalen, 2008), or on sizes at maturity (Stam, de Leemkule, & Ernsting, 1996), we extend these results to asymptotic body lengths.

TO also produces on average larger clutches than HA (Figure S4A), grows faster between 21° and 26°C and reaches slightly larger asymptotic lengths over the whole temperature range (Figure 6). These observations confirm that HA has a slow life history strategy compared to TO (Mallard et al., 2015; Tully & Ferrière, 2008) which remains true under a broad temperature range. Interestingly, we show that the faster lineage also exhibit the stronger TSR response in population. A similar result has been shown in a multi-species comparison (Horne et al., 2015): multivoltine arthropods exhibit a steeper decrease in size with increasing temperature than univoltine ones. We also found that the optimal temperature is higher for HA than TO when looking at reproduction, but not when looking at growth. The significance of this discrepancy remains to be studied.

In populations, the genetic difference in growth rates vanishes and the genetic difference in asymptotic body lengths reverses: adults TO are much smaller than HA over the whole temperature range. This reduction and reversal of genetic differences from one environment to another might result from genetic difference in resource acquisition, resource assimilation or from differences in resource allocation. Indeed, the lineages differ in their ability to invest into reproduction (Figure 2A, B, Tully & Ferrière, 2008). The fast lineage (TO) may require more nutritional resources to grow faster while reaching larger size and reproducing more than the slow one (HA). In *Daphnia*, genetic diversity in resource allocation can generate differences in body size and age at maturity (Hoefnagel et al., 2018). When food becomes scarce in populations due to intraspecific competition, the gap between the resource requirement and its availability may become especially large for TO. The individuals would then be constrained to reduce the resource allocated to growth to favour reproduction which may result in the comparatively large number of juveniles observed in the TO populations (Figure S4B). TO may also have an increased resource intake (which is advantageous when the resource is unlimited) but a lower assimilation efficiency, resulting in smaller average sizes in populations where the resources become limiting. Together, our observations support the idea of genetic variation in the fraction of resource allocated to reproduction and growth (De Jong & van Noordwĳk, 1992). Further work is needed to study whether this genetic variation in resource allocation is linked to a putative variation in resource acquisition and assimilation efficiency and more specifically to examine whether it is advantageous for organism with relatively low assimilation efficiency (TO vs. HA or large vs. small individuals) to slow growth and invest more in reproduction when resources become scarce.

To conclude, our study provides evidence for within-species genetic variation in thermal reaction norms and in plastic responses to intraspecific competition. By providing a quantitative analysis of the way growth rates and asymptotic lengths change according to temperature and density in a population context, we stress the need for untangling the complex interactions between environment and demography. Our results reinforce the idea that the TSR response of ectotherms can be modulated by biotic and abiotic stressors when studied in non-optimal laboratory experiments.

## Acknowledgements

We thank two anonymous referees for their constructive comments that helped us to refine and clarify the manuscript.

We would like to pay tribute to David Claessen’s memory. He left us in July 2018 and we are deeply sorrow for the loss of an unforgettable friend and such a brilliant colleague. We are grateful for his contribution to this work.

## Authors’ contributions

FM, VLB, TT conceived the ideas and designed methodology. FM, VLB, CLC, MA, RP and TT collected the data. FM, VLB, TT analysed the data. FM and TT led the writing of the manuscript. VLB DC and CLC participated to the writing of the manuscript.

All authors contributed critically to the drafts and gave approval for publication.

## Supporting Information

### Supporting methods

#### (i) The Collembola *Folsomia candida* as a model organism

The Collembola *Folsomia candida* is a broadly distributed throughout the world. It is found in habitats such as decaying litter, rotting wood or in caves. It is easy to maintain in the laboratory, and can be bred in isolation or in populations in simplified microcosms with a fine control of temperature, humidity, food provisioning and density (Fountain & Hopkin, 2005). This species is known for its highly flexible phenotypic adjustments when experiencing a sudden change in density or re-source abundance (Tully & Ferrière, 2008). As a parthenogenetic species, multiple individuals sharing the same genotype can easily be bred in different environmental conditions. We worked with two distinct genetic clonal lineages (labelled respectively TO and HA) with contrasted ecological history and bio-demographic strategies (Tully, D’Haese, Richard, & Ferriere, 2006; Tully & Ferrière, 2008; Tully & Lambert, 2011; Mallard et al., 2015): at 21°C at low density, HA individuals have on average a lower reproductive potential and a lower basal mortality than TO individuals and they reach a higher body length. But the ecological natural conditions in which these lineages are adapted are not known.

#### (ii) Rearing of isolated individuals and of populations at different temperatures

Both isolated individuals and populations were kept in standard rearing boxes made of polyethylene vials (diameter 52 mm, height 65 mm) filled with a 30 mm layer of plaster of Paris mixed with 600 mL of Pébéo graphic^®^ Chinese ink to increase visual detectability of light individuals against the dark background (Tully & Ferrière, 2008). Rearing boxes were kept in incubators (Velp FOC 225E, temperature controlled ±0.5°C) at six temperatures (6°, 11°, 16°, 21°, 26° and 29°C) at constant humidity (~100%) and in darkness. Food was provided in the form of small pellets of a mixture of dried yeast and agar in standardised concentration and volume (5000 mL water+80 mg agar+800 mg dried yeast, to produce pellets of 15 μL).

Isolated individuals came from new-borns that were isolated immediately after hatching and fed *ad libitum* during their whole life. We studied between 12 and 20 individuals per lineage and temperature (mean 15.4).

Populations were founded with 5 to 10 adults and were kept at the same temperatures as those for isolated individuals (Figure 1). We were not able to maintain viable populations at 29°C because at this temperature the collembola do not reproduce (Figure 2A). Between four and twelve populations were started for each lineage at each temperature (Figure S2).

#### (iii) Measuring densities of large and small individuals

We used the option *region = TRUE* of the *STdiag.measure* function to extract a region of the plot and get from it the number of individuals and the length of each individual during the periods of measurements. More specifically, for measurements of asymptotic body lengths we estimated the mean density of large individuals (which we called “adults”) and small ones (also called “juveniles”) at the time of each measurement as the mean density during a five weeks period which includes the week of the measurement, three weeks before and the week after the measurement (Figure 1, Figure 2B). We used five weeks periods to get average measurements of the density integrating fluctuations of density that could occur during the period of stabilization of the mean body length of the studied cohorts. For growth rate measurements, the density of large and small collembola were estimated as the mean densities over the period of time on which the growth rate has been estimated.

We used the number of large individuals (on a log scale) to quantify the relationship between temperature and “adult” density (Figure 2B) and to further measure the strength of competition by quantifying the dependence of growth rate and asymptotic length to adult density (Figure 4). We used the logarithm of adult density as a proxy of the population state since adults represent on average 90% of the total “biosurface’ (total surface of collembola estimated with our pictures) in our populations and since we knew from previous experiments that large individuals play a central role in the density dependent effects by outcompeting small individuals in resource acquisition (Le Bourlot, 2014; Le Bourlot et al., 2014).

Note that when two modes of large individuals could be identified (HA populations at 21°C, Figure S2), we only took into account the number of very large individuals (>1.5 mm) because we know from the previous experiments mentioned above that the large collembola dominates not only the juveniles but also the smaller adults.

#### (iv) Measuring the mean growth rates and asymptotic body lengths in populations

##### Cohort growth rates

To measure the growth rates of the cohorts on the structure-time diagrams (Figure S2), we adjusted a strait line rather than a growth curve because many cohorts that are well distinguishable during their period of maximum growth are less well distinguishable before and after this period especially when they merge with other cohorts.

##### Asymptotic body length

We found no other reliable way to automatically measure the cohorts′ asymptotic body lengths due to the complexity of the structure-time diagrams, especially when several cohorts fuse or when several modes of large collembola coexist (See the example in Figure 1 and some populations such as TO_21_r2 in Figure S2).

### Supporting figures

**Figure S1:**
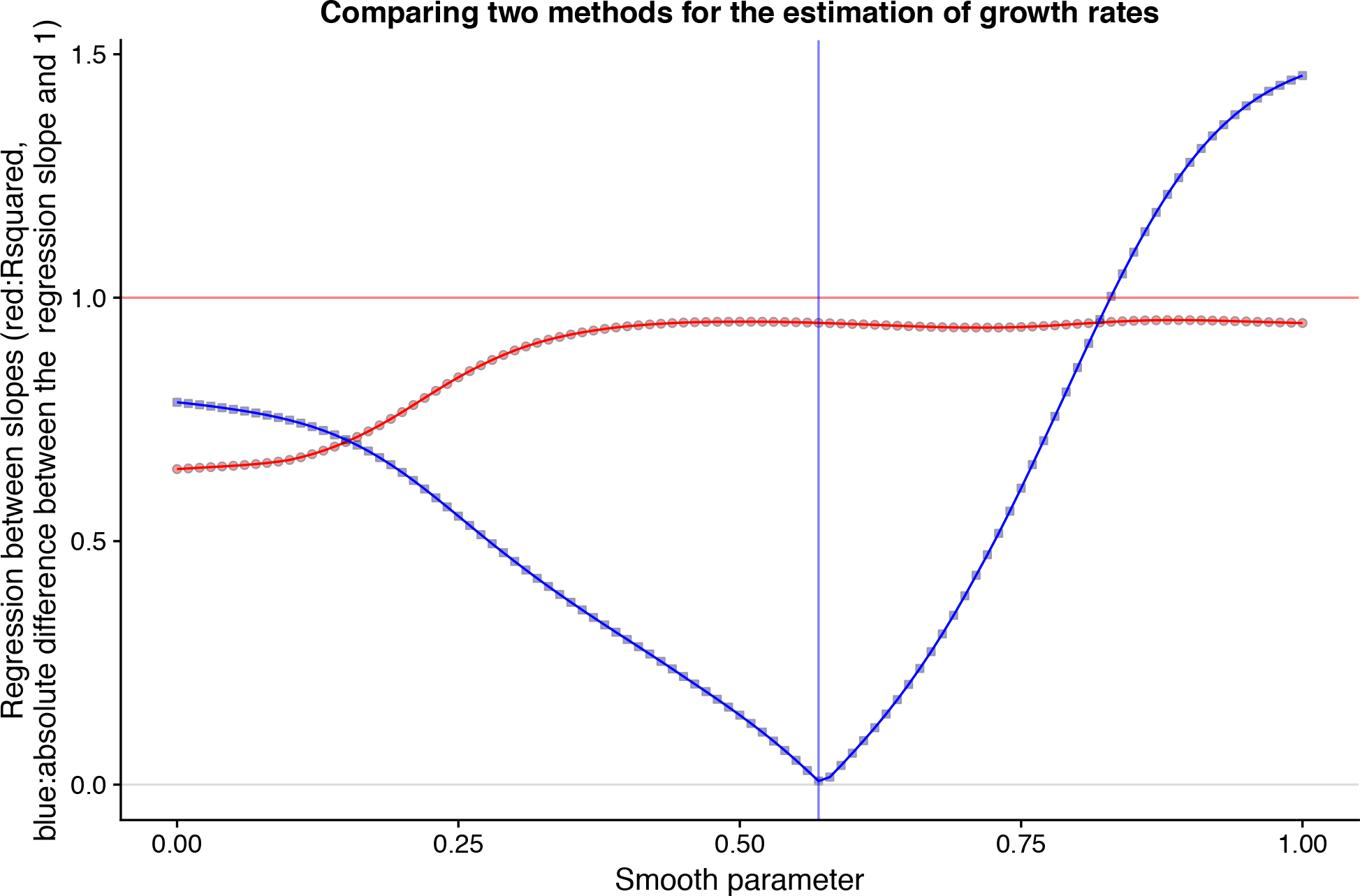
Comparing methods to measure growth rates. To verify that the method used to measure the growth rates in populations provides estimates that we can safely and reliably compare to the ones made on isolated individuals, we have studied in details 14 cohorts than can be easily identified on the plots and we have extracted about 17 points on each of these cohort’s growth trajectories. We used this dataset to compare growth rates estimated by eye on these cohorts with growth rates estimated by fitting a parametric growth curve. More specifically, we fit smoothed spline growth curves on these 14 cohorts, using the *gcFitSpline* function from the *grofit* package (Type=Gompertz). These fits were done recursively with different values of the *smooth.gc* parameter (between 0 and 1) and for each of these values, we measured the R-squared and the slope of a regression between the growth parameters estimated by eye and the growth parameters estimated from the smoothed splines. The figure shows how varying the smooth parameter of the spline curves modifies the quality of the regression between the two types of measurements. The red curve shows the R-squared for different values of the smooth parameters. The blue curve quantifies how far the regression slope is from a slope of one. The R-squared reach values higher than 0.95 for *smooth.gc* parameter above 0.45 and a *smooth.gc* parameter of 0.57 minimizes the difference between the estimation of the slope (0.99) and one. Thus, to minimize potential bias between the two methods, we choose to use a smooth parameter of 0.57 to fit the growth curves of the isolated individuals.

**Figure S2:**
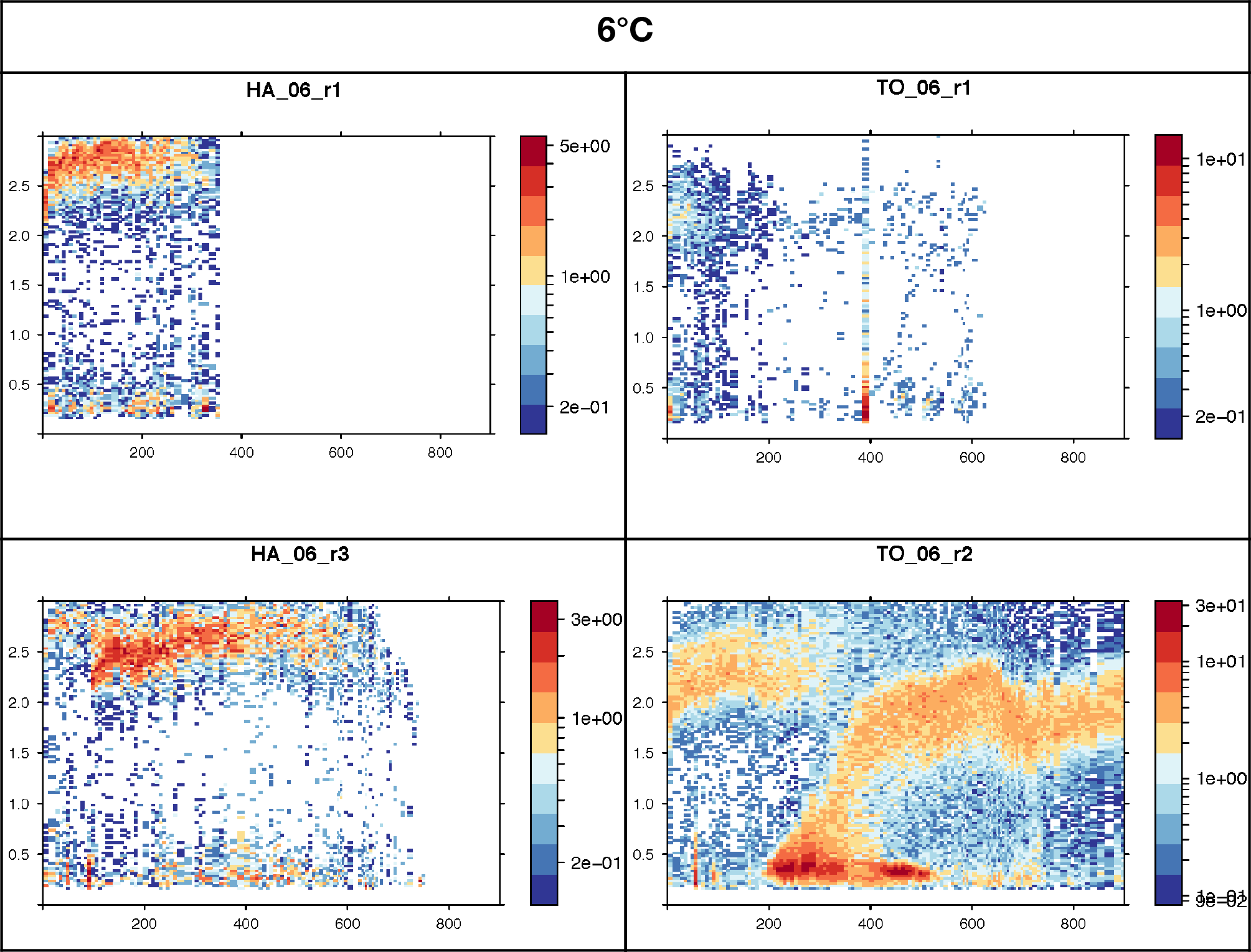
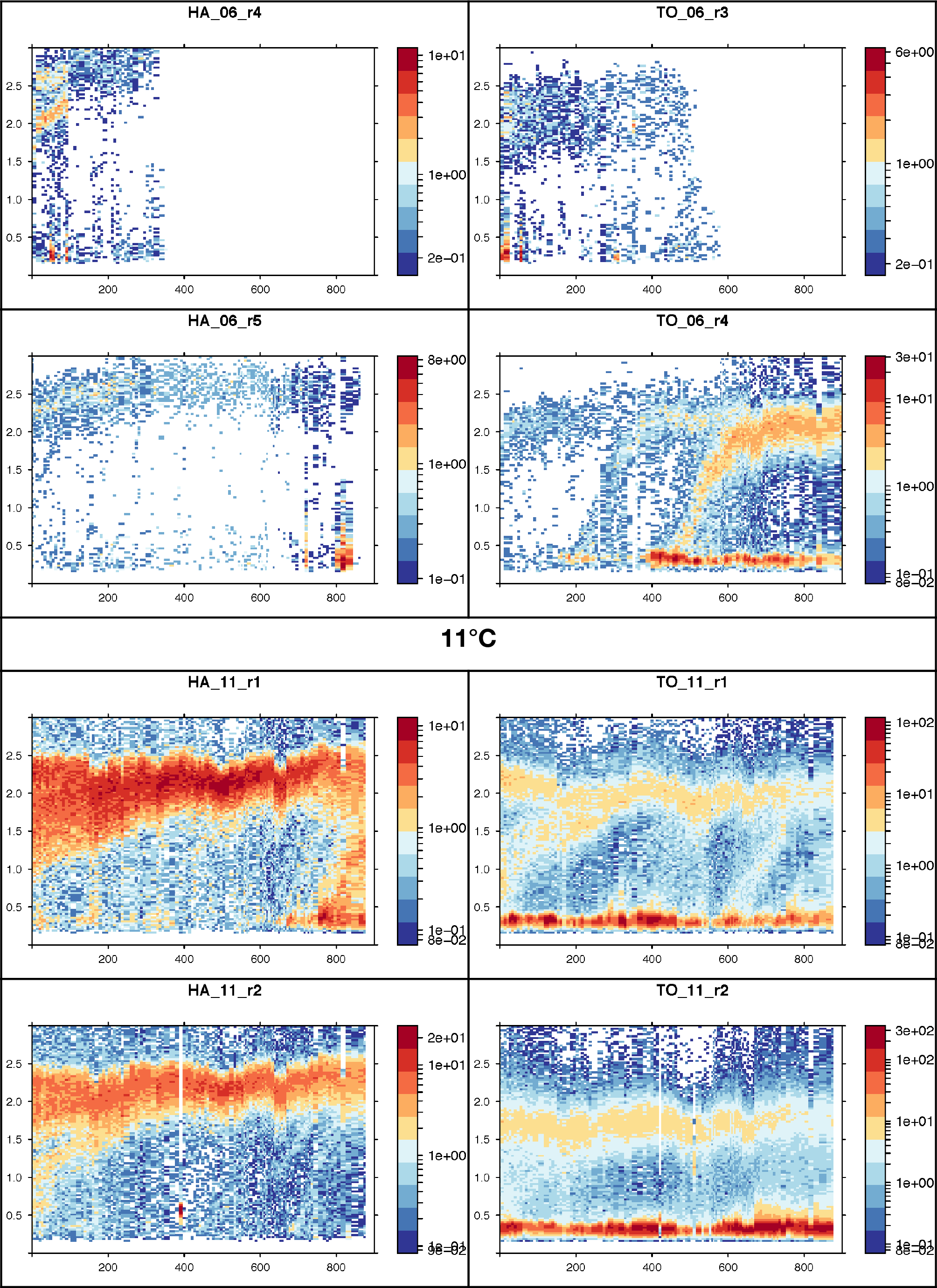
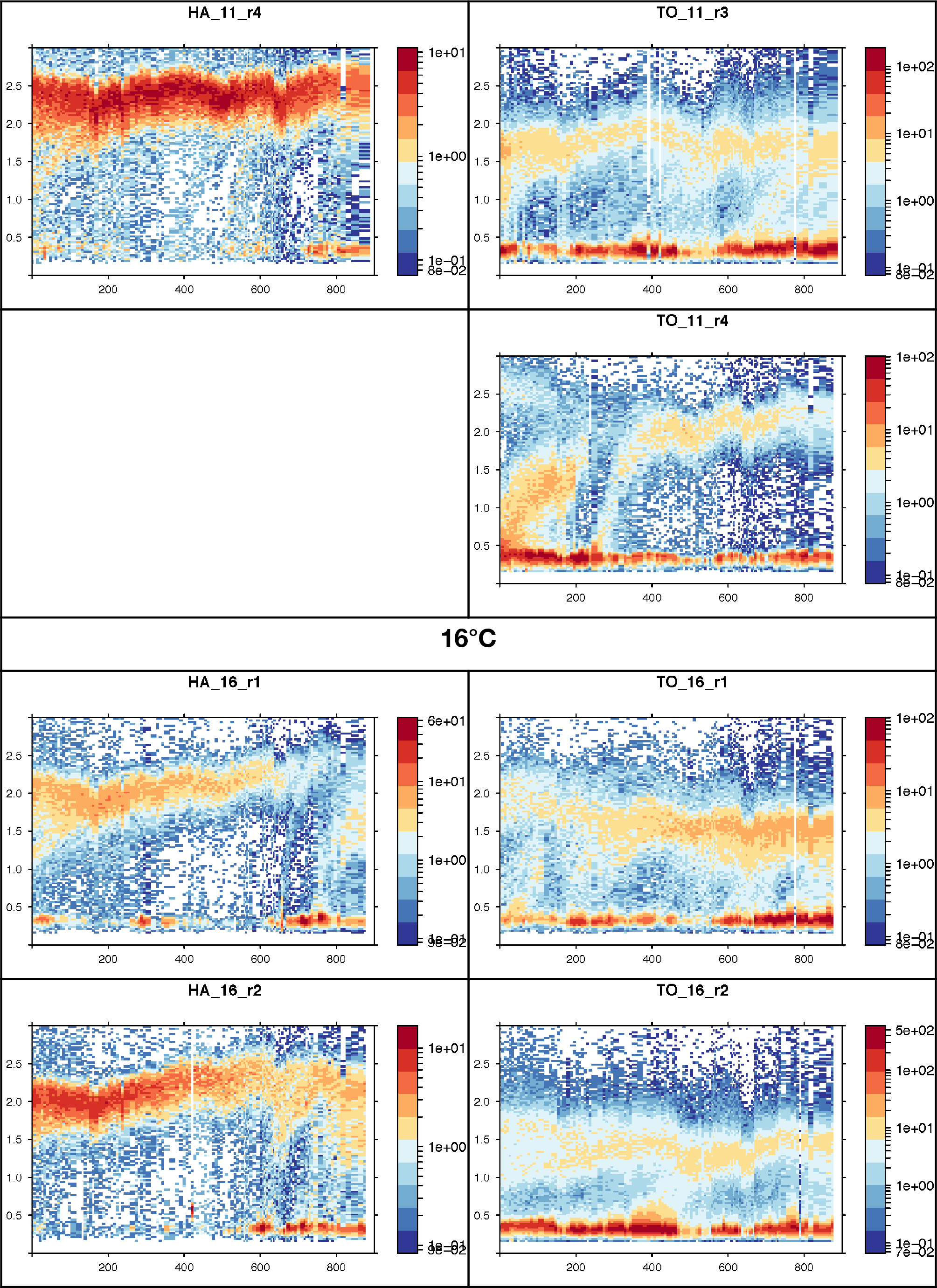
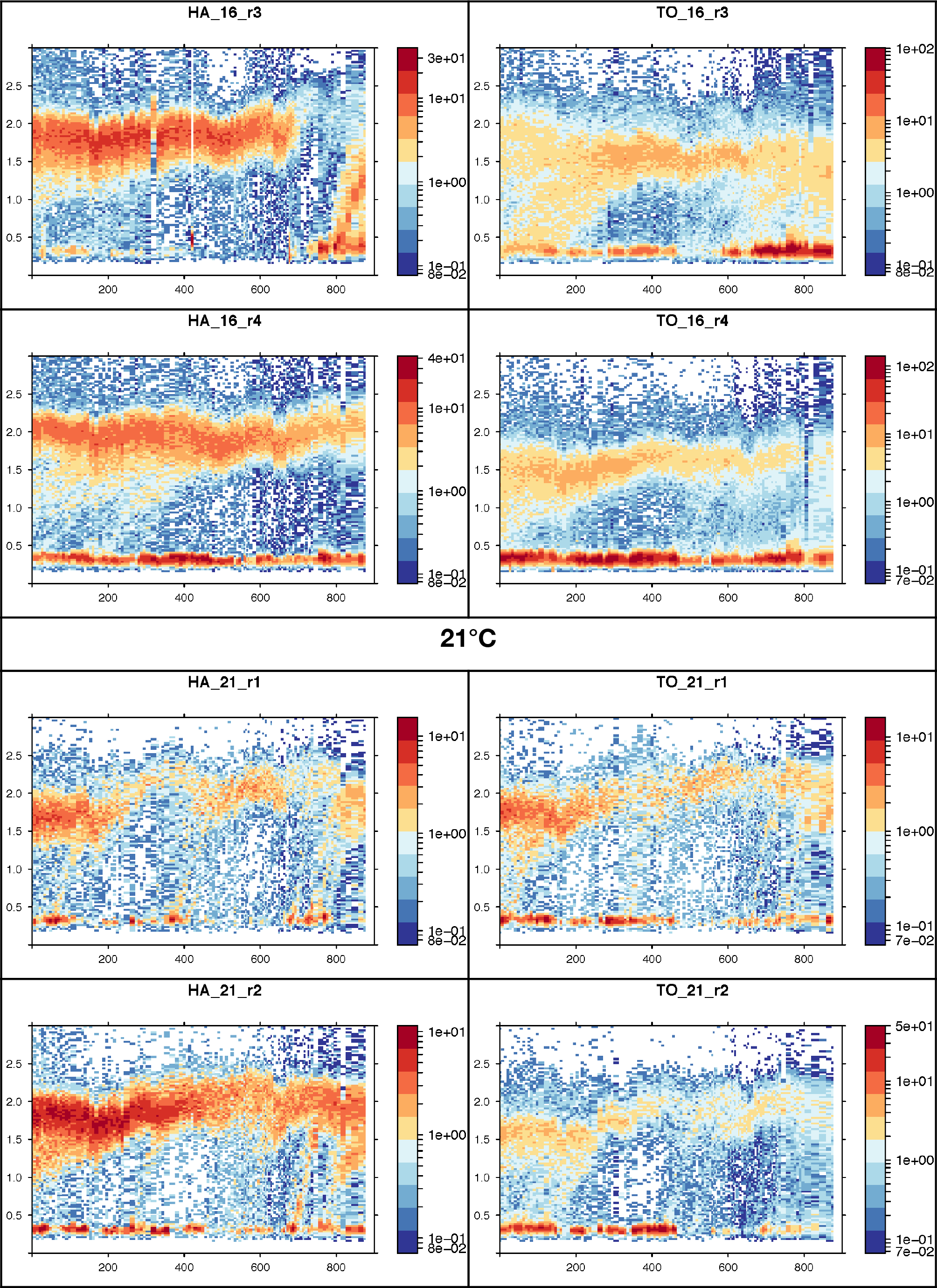
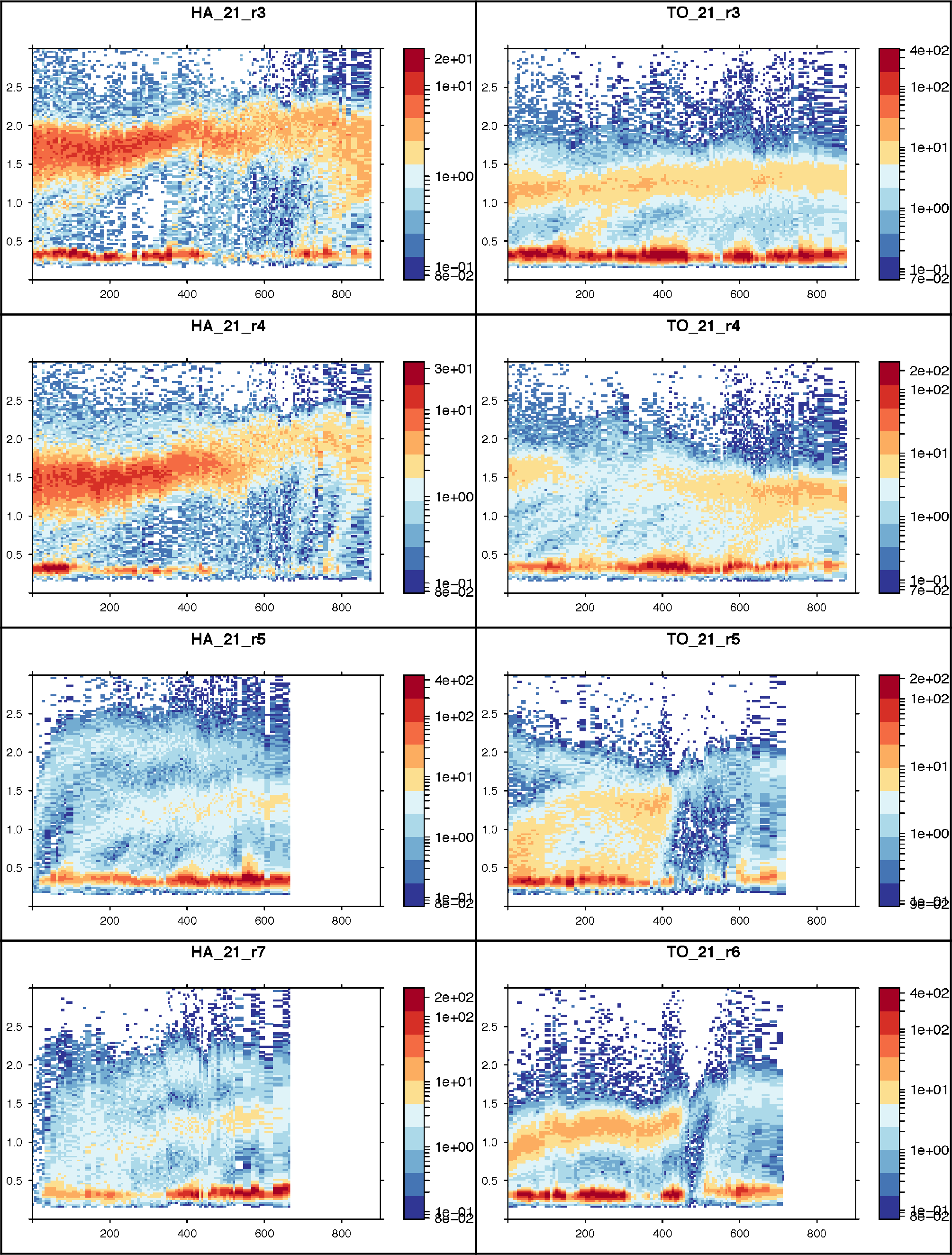
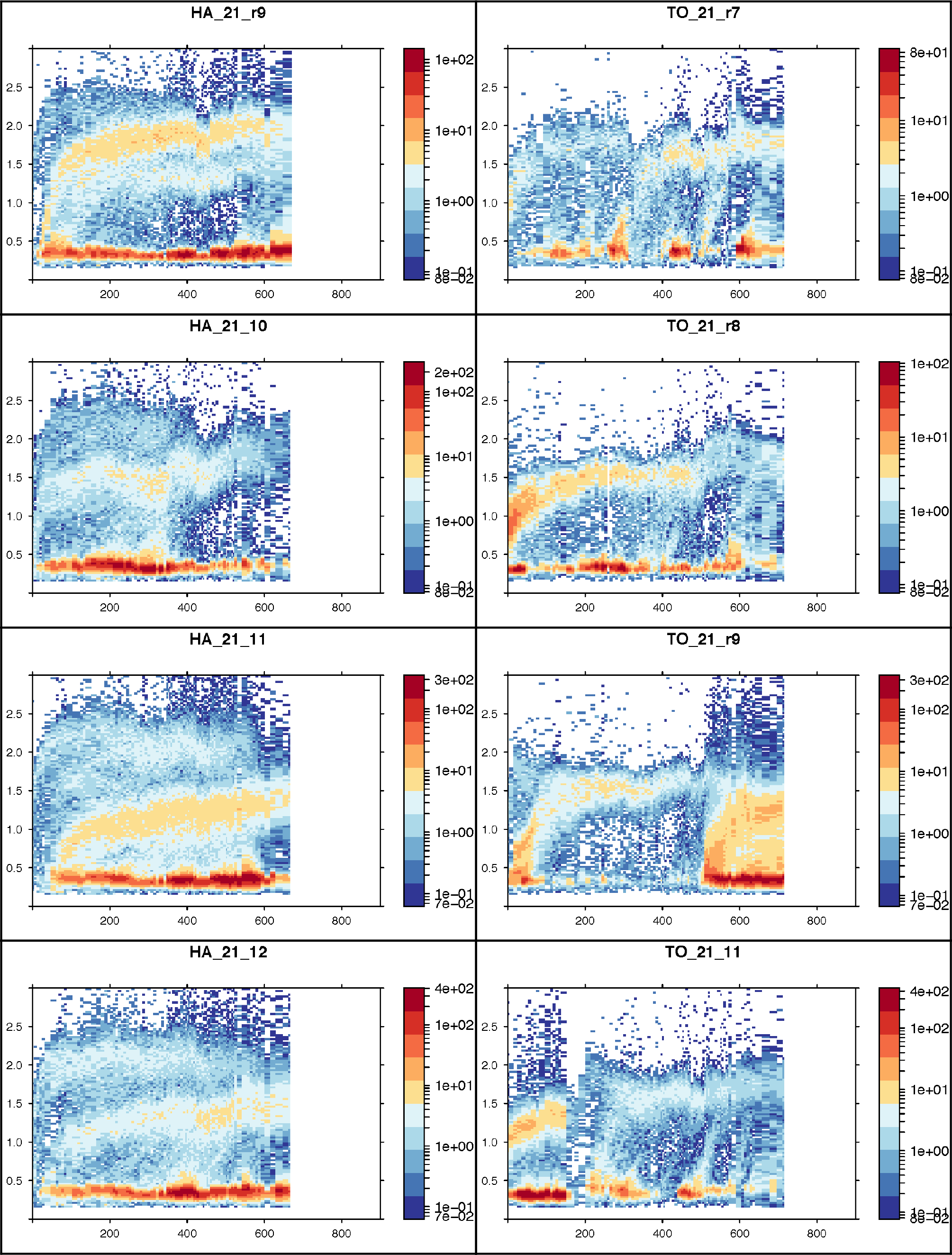
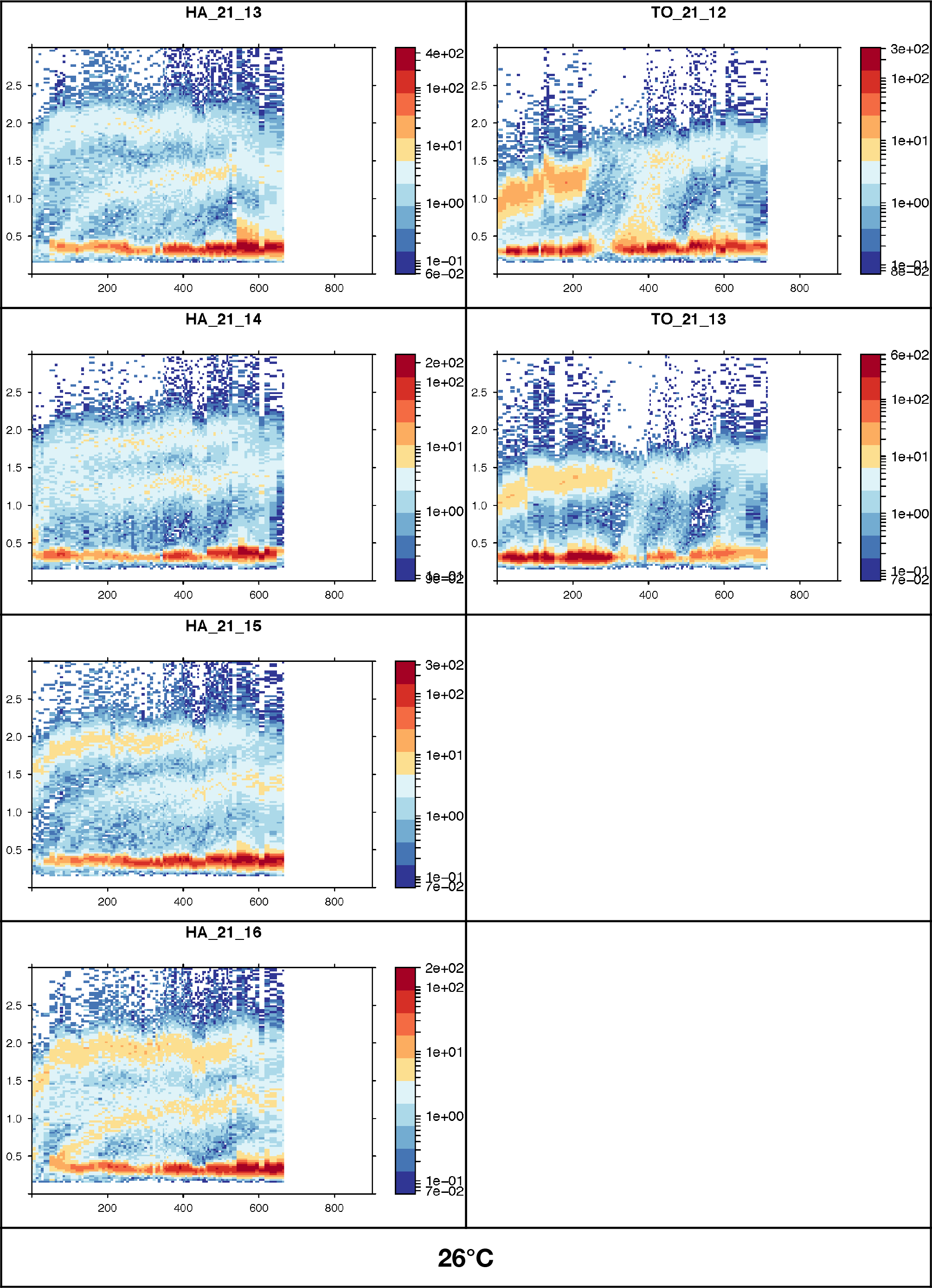
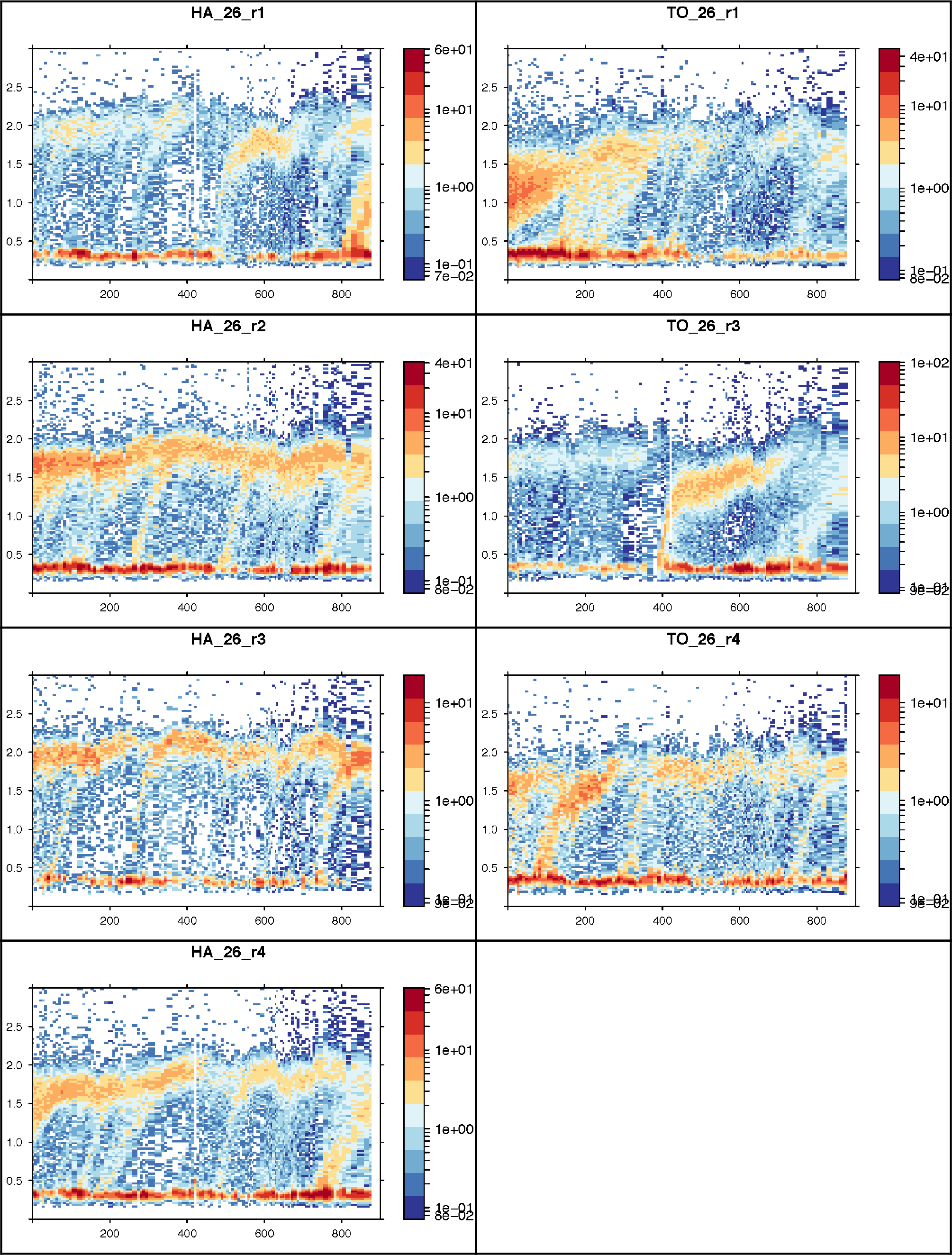
Structure time diagrams of 56 populations from two clonal lineages of Collembola (HA on the left side and TO on the right side) raised at five different temperatures (6, 11, 16, 21 and 26°C). The population labelled « HA_06_r1 » for example is the first population (replicate 1) of HA individuals raised at 6°C. Some populations became extinct and their dynamics are not plotted (ex. HA_11_r3). These diagrams display the temporal dynamics of the populations’ size-structure: for each time (x-axis, number of days) and size class (y-axis, body length in mm) coordinate, a colour rectangle is plotted whose hue refers to the number of individuals (on a log scale, scale on the right side of each plot). We used these diagrams to estimates the growth rate and asymptotic body length of cohorts. All these measurements were done using the function *STdiag.measure* from the R library *STdiag* that we specifically developed for this purpose (Le Bourlot V et al., 2015). This function is designed to interact with the structure-time diagram in order to obtain some quantitative measurements such as growth rate or average length of a group of individuals by clicking directly on the diagram. Note that a difficulty comes when measuring the asymptotic length from the remarkable plasticity of the springtails. They can resume growth or even shrink when the environmental conditions change (the density and structure of the populations change). To deal with this, we chose to focus on the mean length reached by a cohort right after it has stopped or significantly slowed down its growth even if the cohort can resume growth after a while when the population density changes for instance.

**Figure S3:**
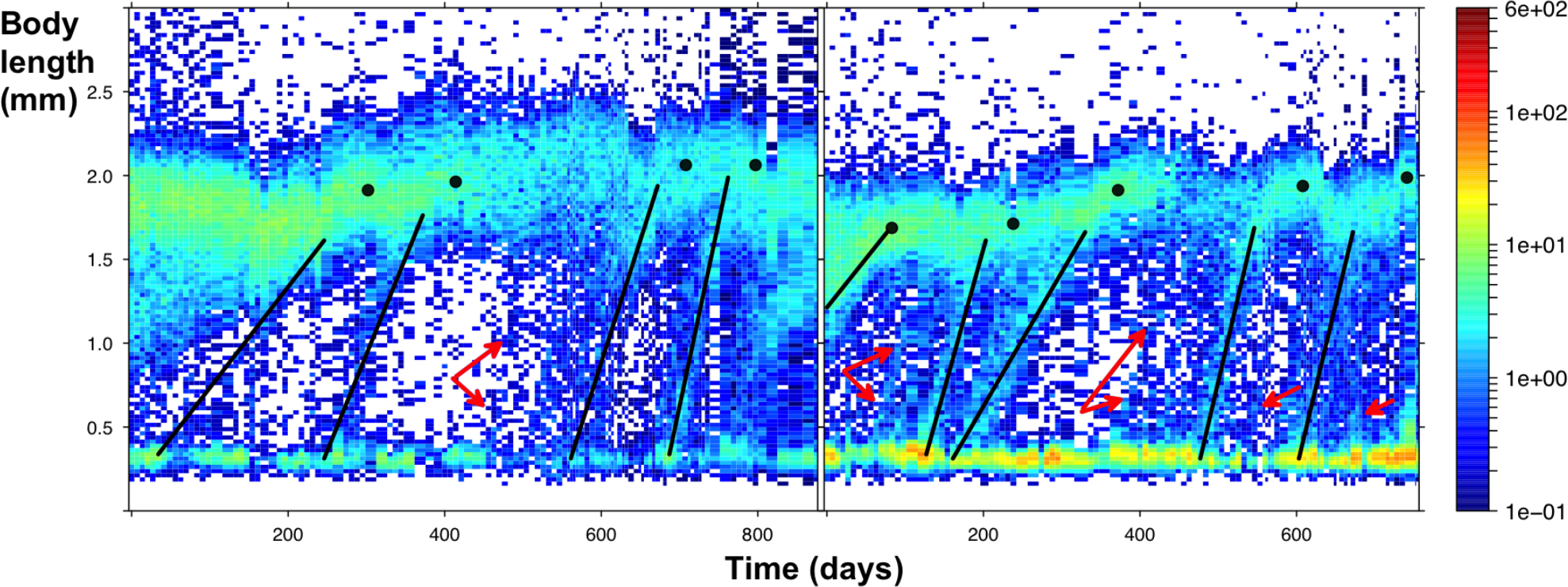
Two populations where different cohorts of juvenile recruiting in adults are visible. Black segments are fitted on cohorts that are sufficiently contrasted to be used to do the growth measurements. Red arrows point toward other cohorts that are less distinguishable and that were not used to estimate their growth rates. The black dots show the measurements of asymptotic body lengths.

**Figure S4:**
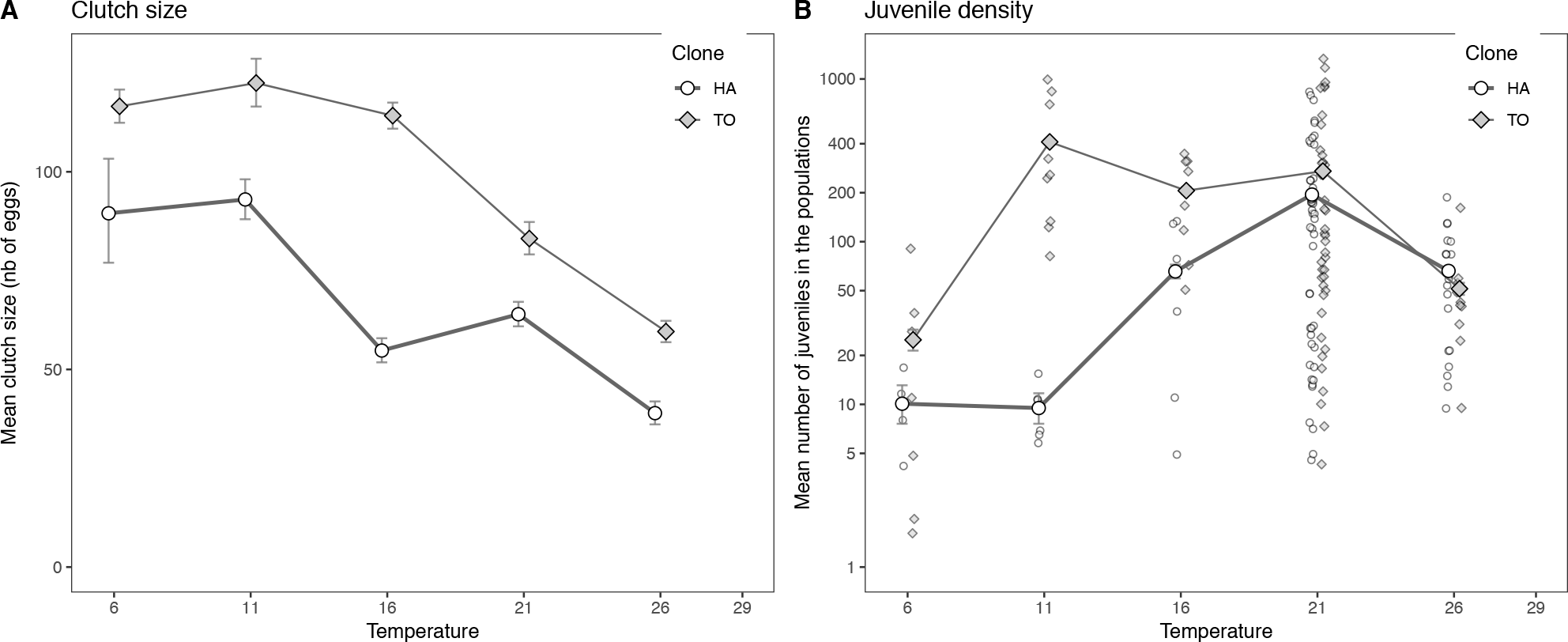
The effects of temperature on the clutch size produced by isolated individuals (A) and on juvenile density in populations (B) for lineages HA (open circle) and TO (grey diamonds). Panel A presents mean clutch sizes (mean numbers of eggs per clutch measured on 336 clutches, +/− 95% confidence interval) as a function of temperature. The TO lineage lay on average larger clutches than HA. The juvenile density in populations is plotted as a function of temperature on panel B (see methods for details). On average, there are more juveniles in TO populations than in HA ones.

**Figure S5:**
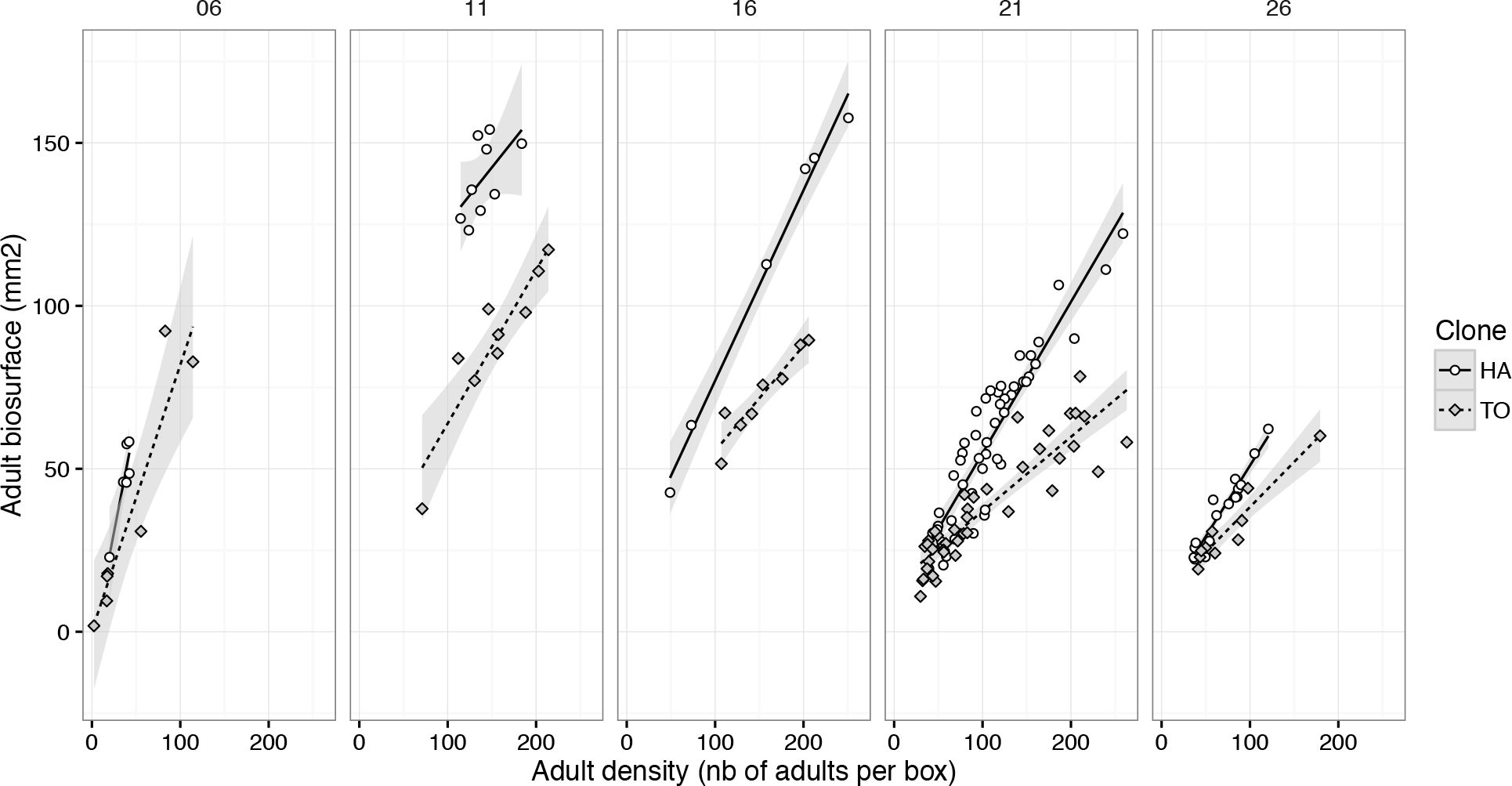
Relation between the adult biosurface (sum of the surface of all the adults in a box in mm2) and the adult density (number of adults per container) for the two lineages and at the five temperatures. For the same adult density, the biosurface of HA is higher than TO because on average in populations adults HA are larger than TO.

**Figure S6:**
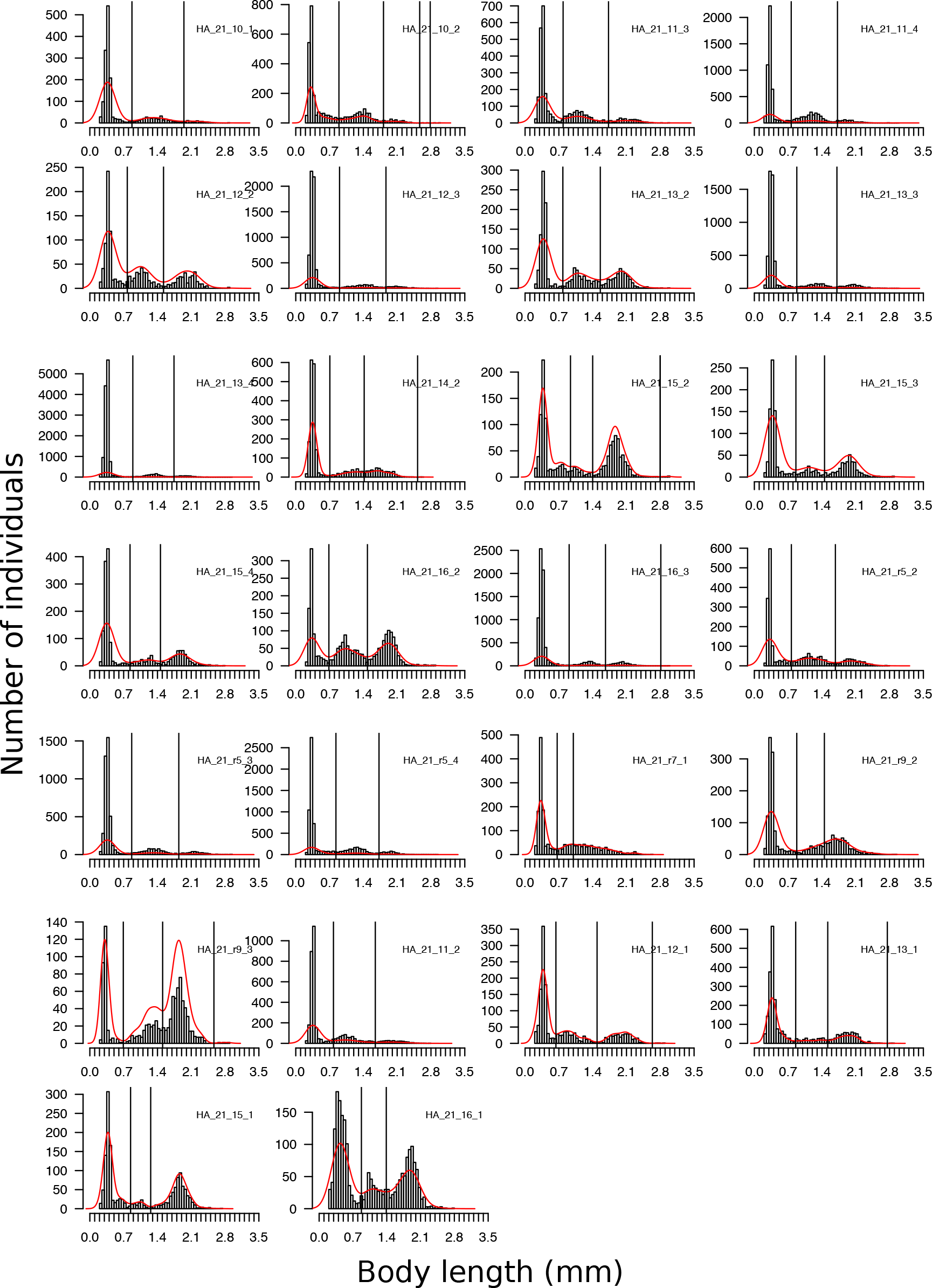
Trimodal body length distributions observed in some of HA populations raised at 21°C when the mean asymptotic size attained by some cohorts was measured (Figure S1). A kernel density estimation (red curve) has been fitted to the raw distribution (black histogram). The extremes of these red curves were used to automatically split each size distribution into three size classes separated here by black vertical lines.

**Figure S7:**
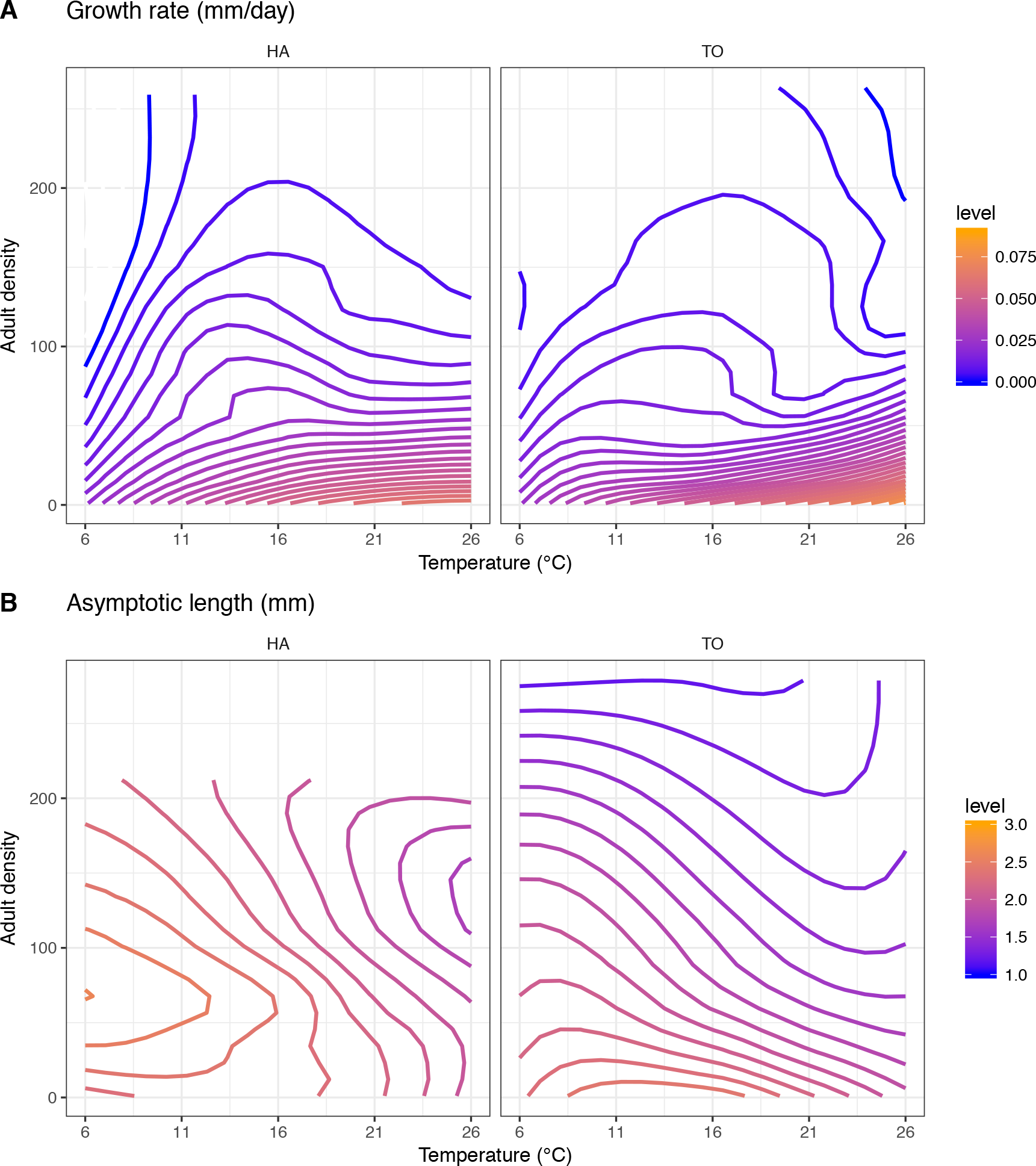
The visual display of the joint effect of temperature and density on growth rates and asymptotic body lengths using data measured on isolated individuals (Adult density of 1) and in populations. Panel A shows the positive effect of warming on growth rate and the additive negative effect of density (competition) on growth. One can also see that above a certain density, the isoclines become more or less horizontal. This means that, as the density increases, the temperature has less and less effects on the growth rates, the main variable that limits growth being the competition. Panel B shows that for the very low densities, the asymptotic length decreases with increasing temperature for both lineages. For TO, the interspecific competition has a negative effect on asymptotic length with an additive effect of temperature that remains on almost the whole range of conditions explored. For HA, the density has first a positive effect on asymptotic length: when the adult density increases between 1 and about 50 adults per container, the collembola tend to reach a longer asymptotic body length. But when the density continues to increase, its effect becomes negative (the isoclines are bended and their slopes change, B).

### Supporting tables

**Table S1:**
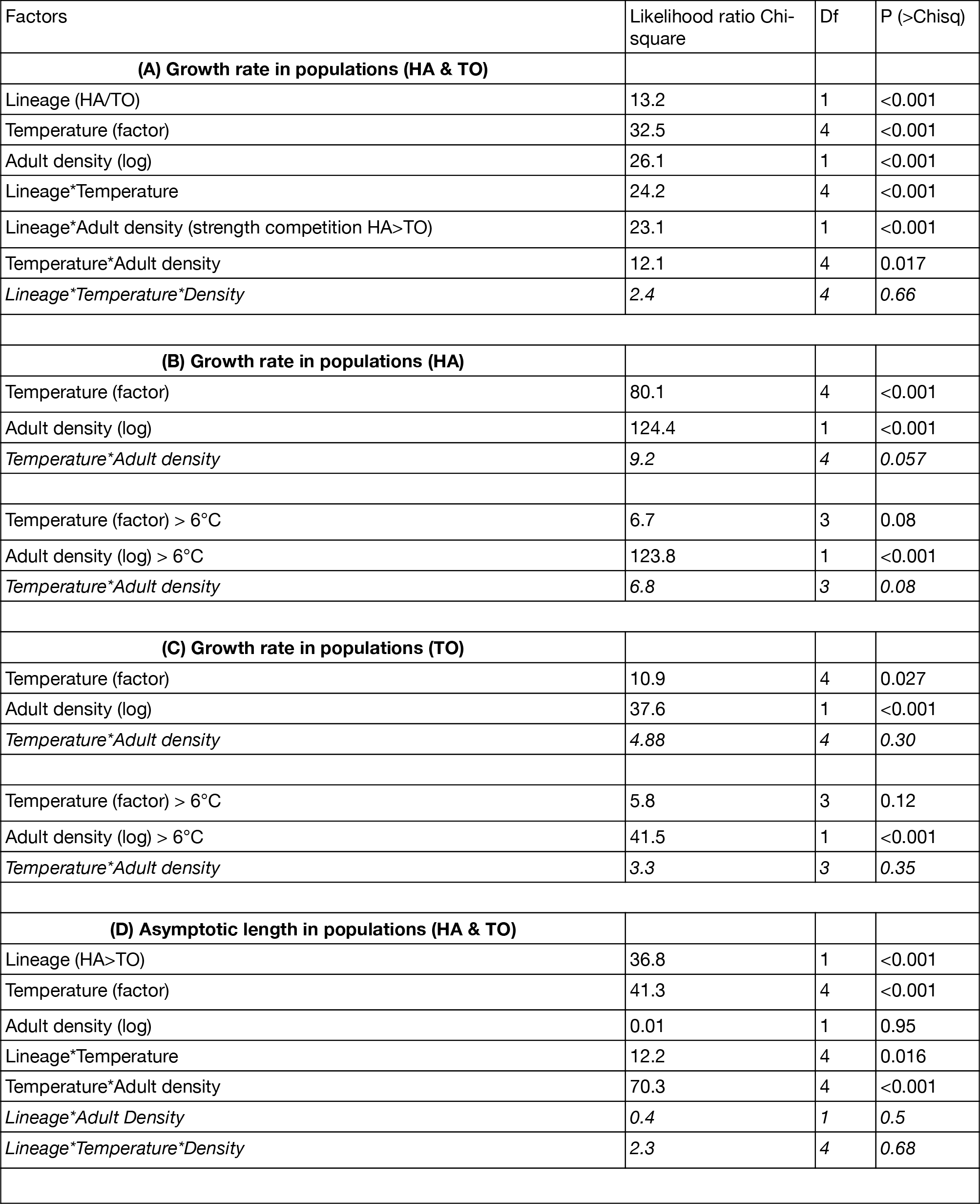
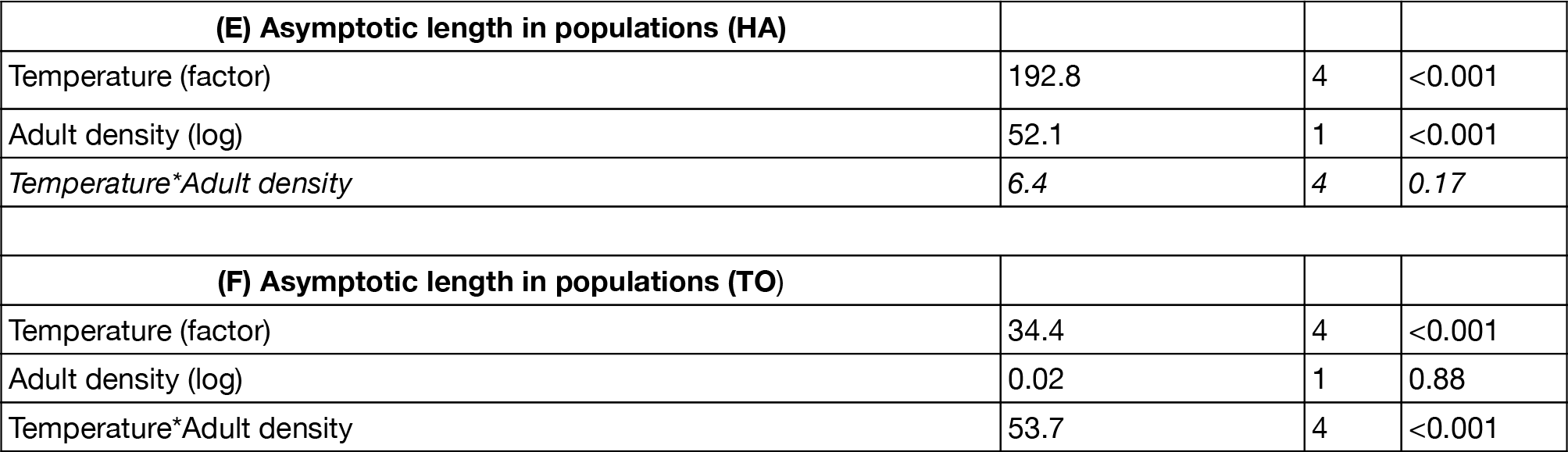
Best selected linear models describing the effect of *adult density* (log scale), *temperature* (as a factor) and *lineage* identity (HA and TO) on the *growth rate* (models A-C) and *asymptotic body length* (models D-F) measured in populations (Figure 4). Non-significant complex interactions that have been dropped from the initial full models are italicized. Effects and their interactions are tested with likelihood ratio tests using type 3 anova (*Anova* function from the *car* package). The main models (A, D) are associated with sub-models (B-C, E-F) for each of the two lineages.

**Table S2:**
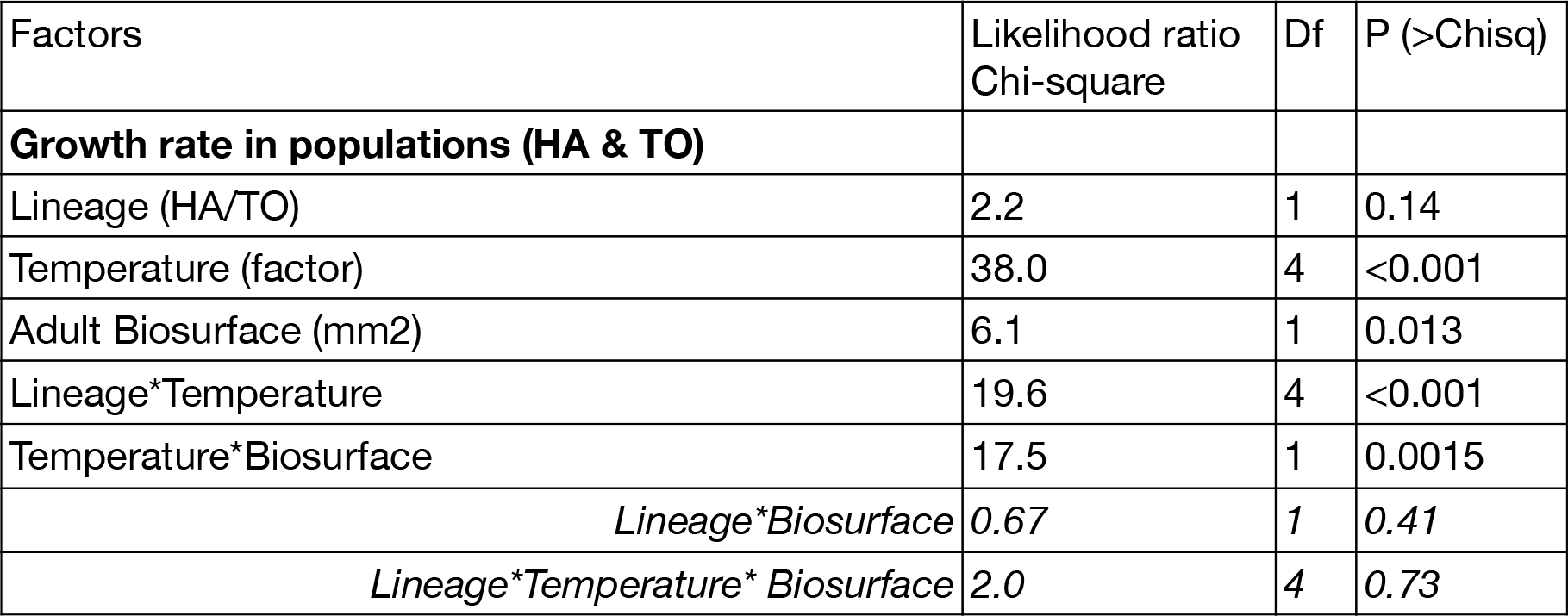
Model including the adult biosurface (sum of the surface of adults in populations) as a covariable rather than number of adults to explain the observed variations of cohort growth rates. When the total surface of adults (biosurface) is taken into account rather than their number, the previously observed difference between the two lineages (significant interaction between lineage and density in model A in Table 3 suggesting that the strength of competition in higher for HA) vanishes. This shows that the strength of competition is the same between the two lineages when controlling for adult size.

**Table S3:** This table gathers the estimates (mean and 95% rate) and B (asymptotic body length) from Table 1. The predicted asymptotic body length of the clone HA in the population raised at 6°C is 2.54mm. At 21°C, the predicted asymptotic size of isolated TO is 2.3 mm (2.54-0.89-0.46-0.37+0.43+1.03+0.64-0.62=2.3).

|  | Growth rate (µm/day) | Asymptotic body length (mm) |
| --- | --- | --- |
| Intercept (6°, HA, population) | 8 [2.7 ; 13.3] | 2.54 [2.31 ; 2.76] |
| 11° | 1.9 [-4.9 ; 8.7] | -0.19 [-0.47 ; 0.09] |
| 16° | 3.7 [-3.7 ; 11.2] | -0.44 [-0.75 ; -0.14] |
| 21° | 3.8 [-1.8 ; 9.4] | -0.89 [-1.13 ; -0.66] |
| 26° | 13.2 [7.1 ; 19.2] | -0.62 [-0.88 ; -0.37] |
| TO | 5.6 [-1.6 ; 12.8] | -0.46 [-0.75 ; -0.16] |
| Isolated individual | 15.7 [5.2 ; 26.3] | -0.37 [-0.79 ; 0.05] |
| TO 11° | -8.6 [-18 ; 0.9] | -0.1 [-0.48 ; 0.28] |
| TO 16° | -10 [-20.1 ; 0] | -0.07 [-0.47 ; 0.33] |
| TO 21° | -6 [-13.7 ; 1.7] | 0.43 [0.12 ; 0.74] |
| TO 26° | -11.4 [-20.2 ; -2.6] | 0.21 [-0.14 ; 0.57] |
| Isolated individual 11° | 10.9 [-1.9 ; 23.7] | 0.32 [-0.18 ; 0.83] |
| Isolated individual 16° | 24.6 [11.5 ; 37.8] | 0.61 [0.09 ; 1.13] |
| Isolated individual 21° | 38.1 [25.6 ; 50.7] | 1.03 [0.54 ; 1.53] |
| Isolated individual 26 | 27.7 [15.3 ; 40.1] | 0.35 [-0.14 ; 0.84] |
| TO Isolated | -5.6 [-18.3 ; 7] | 0.64 [0.14 ; 1.14] |
| TO Isolated individual 11° | 14.5 [-1.9 ; 31] | 0.12 [-0.53 ; 0.77] |
| TO Isolated individual 16° | 9.1 [-7.1 ; 25.3] | 0.01 [-0.63 ; 0.65] |
| TO Isolated individual 21° | 11.8 [-5.3 ; 28.9] | -0.62 [-1.29 ; 0.05] |
| TO Isolated individual 26° | 29 [13,5 ; 44,6] | -0,4 [-1,02 ; 0,21] |

